# Transcranial theta-burst stimulation of primary sensory cortex attenuates somatosensory threat memory in humans

**DOI:** 10.1101/2021.06.09.447685

**Authors:** Karita E. Ojala, Matthias Staib, Samuel Gerster, Christian C. Ruff, Dominik R. Bach

**Affiliations:** Computational Psychiatry Research, Department of Psychiatry, Psychotherapy and Psychosomatics, Psychiatric Hospital, University of Zurich, Switzerland; Neuroscience Centre Zurich, University of Zurich, Switzerland; Zurich Center for Neuroeconomics (ZNE), Department of Economics, University of Zurich, Switzerland; Wellcome Centre for Human Neuroimaging and Max-Planck UCL Centre for Computational Psychiatry and Ageing Research, University College London, UK

## Abstract

Sensory cortices are required for learning to discriminate complex stimuli that predict threat from those that predict safety in rodents. Yet, sensory cortices may not be needed to learn threat associations to simple stimuli. It is unknown whether these findings apply in humans. Here, we investigated the role of primary sensory cortex in discriminative threat conditioning with simple and complex somatosensory conditioned stimuli (CS) in healthy humans. Immediately before conditioning, participants received continuous theta-burst transcranial magnetic stimulation (cTBS) to primary somatosensory cortex either in the CS-contralateral or CS-ipsilateral hemisphere. After overnight consolidation, threat memory was attenuated in the contralateral compared to the ipsilateral group, as indicated by reduced startle eye-blink potentiation. There was no evidence for a difference between simple and complex stimuli, or that CS identification or conditioning was affected, suggesting a stronger effect of cTBS on consolidation than on initial stimulus processing. We propose that non-invasive stimulation of sensory cortex may provide a new avenue for interfering with threat memories in humans.

## Introduction

Discriminating sensory predictors of threat and safety is important for adaptive functioning in the environment and may be disrupted in anxiety and fear disorders^1,2^. However, the role of sensory cortices in learning from threat remains debated. This is an important question not only to understand the neural macrocircuits that underlie the computational machinery of associative learning^3,4^ but also to gauge the potential utility of clinical interventions with non-invasive brain stimulation for which sensory cortices could constitute an accessible entry point^5,6^. It is well-known from studies in rodents that acquisition of threat memory depends fundamentally on synaptic plasticity in basolateral and centromedial amygdala^7,8^. The amygdala receives information about conditioned stimuli (CS) and unconditioned stimuli (US) directly via sensory thalamus^9^ and, in the case of auditory threat conditioning, via an additional indirect pathway through auditory cortex^9,10^. Similar cortico-amygdala projections exist for all sensory modalities across species^9,11^, and indeed there is evidence that secondary auditory, visual and olfactory cortices are involved in rodent threat conditioning^12^ and show post-learning plasticity^13^.

Yet, the precise role of sensory cortices may depend on the complexity of the CS and remains controversial for some types of threat associations. That is, acquisition of non-discriminative threat conditioning with an artificial simple frequency tone as CS may not depend crucially on auditory cortex^12,14–20^ (although note^19,21^). The case is less clear for discriminative learning with two simple CS: optogenetic inactivation of auditory cortex left learning unimpaired in one study^22^, but muscimol inactivation of auditory cortex^23^ and electrolytic lesions of the cortex-projecting ventral division of the auditory medial geniculate nucleus^24^ impaired learning in other studies. In contrast, auditory cortex may have a key role in associating naturalistic or complex auditory CS (e.g., frequency sweeps) with threat US^22,25–27^ (although note^28^). Specifically, there is evidence that disrupting neural transmission in auditory cortex during learning impairs threat memory acquisition of complex stimuli^22,26^ and blocking protein synthesis impairs posts-learning consolidation but leaves acquisition unaffected^27^. Finally, disrupting neural transmission in auditory cortex during cue presentation in a post-conditioning test impairs complex discriminative but not non-discriminative threat memory retrieval^20^.

In human studies, threat-predictive CS+ and safe CS− elicit differential evoked-potential amplitude, oscillatory synchrony, and univariate fMRI BOLD amplitude in visual cortex^29–31^, multivariate BOLD patterns in olfactory cortex^32^, and univariate BOLD amplitude as well as multivariate patterns in the auditory cortex, with no difference between simple and complex CS^33–35^. Recent evidence has also shown that the visual cortex is involved in long-term threat memory in humans^36^. However, the precise function of these areas in the acquisition and retention of threat memories remains unresolved. In particular, it is unclear whether activity in these areas is in fact required for threat memoires to be formed and/or consolidated.

Here, we sought to elucidate the role of primary sensory cortex for discriminative conditioning from simple CS, as well as to translate established rodent findings on complex CS to humans. To this end, we used continuous theta-burst transcranial magnetic stimulation (cTBS) as a technique that can transiently downregulate cortical excitability and synaptic plasticity in the targeted area^37^. As a model sensory system, we selected the somatosensory modality, since primary somatosensory cortex (S1) is well accessible to cTBS and since stimulus representation is strictly contralateral, unlike in the auditory system or in foveal vision^38^. Somatosensory threat conditioning in humans has been demonstrated previously^39,40^, including a study from our own laboratory with the same CS as in the present work^41^. To examine the role of complex stimulus features, we used simple CS defined by stimulus location (index or middle finger), and complex CS defined by a temporal pattern of alternating locations (Fig. 1). S1 was localized individually with functional magnetic resonance imaging (fMRI).

**Figure 1.**
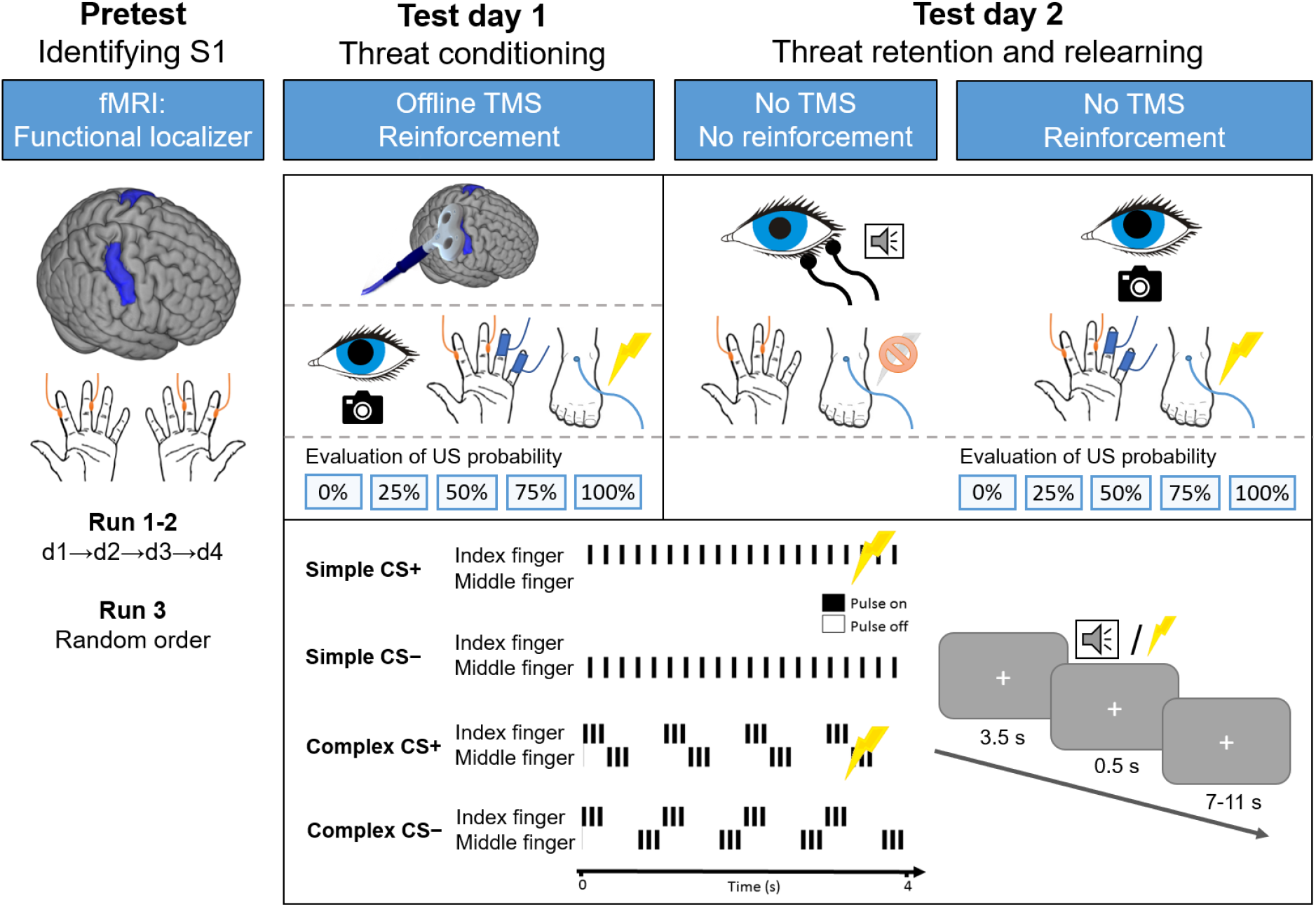
Structure of the study and the experimental paradigm. First, participants underwent fMRI and somatosensory stimulation to functionally localize the primary somatosensory cortex (S1) for the second digits of index and middle fingers (digits d1: left middle finger, d2: left index finger, d3: right index finger, d4: right middle finger). On the first test day, participants received continuous theta-burst TMS targeted on S1. Experimental group received cTBS on the right hemisphere, contralateral to the somatosensory CS stimulation on the left hand. Control group received cTBS on the ipsilateral hemisphere. Immediately after cTBS, participants underwent threat conditioning with aversive electric shock US with a 50% reinforcement rate. US was delivered to the foot ipsilateral to the cTBS site. Simple CS were electric pulses to either index or middle finger of the left hand whereas complex CS were pulses alternating between fingers. Simple and complex CS blocks alternated in all sessions. Each CS lasted for 4 s and 50% of CS+ trials co-terminated with the 0.5-s US, whereas CS- trials never co-terminated with the US. On each trial, participants saw a fixation cross in the center of the screen and were asked to press a button as fast as possible to indicate which of the two possible stimulus patterns they perceived. Pupil size and skin conductance from the ring and little fingers were recorded during conditioning. On the next day, participants came back for a test of threat memory retention and a relearning session without cTBS. Retention session was conducted under extinction (no US) and eye-blink reflex to an acoustic startle probe was recorded. Relearning was conducted afterwards with 50% reinforcement. Pupil size, and skin conductance responses were recorded also during relearning. At the end of the conditioning and relearning sessions, participants rated each CS for how likely they thought it had co-terminated with the US during the experiment.

We expected reduced threat memory retention after overnight memory consolidation when cTBS had been applied to stimulus-contralateral sensory cortex (experimental group), as compared to the stimulus-ipsilateral cortex (control group). We measured threat memory retention by potentiation of the startle eye-blink reflex to CS+ versus CS-. Studies in rodents suggest that cTBS exerts effects at multiple timescales: It can elicit action potentials immediately during stimulation, lead to a period of reduced cortical excitability in the direct aftermath of the stimulation, and induce longer-term changes in learning and memory at different timescales through synaptic long-term depression (LTD) as well as by modulating neurotransmitters, neural growth factors and gene expression^42,43^. In humans, there is evidence that the inhibitory, LTD-like effect of cTBS^37^ may be mediated by effects on both GABAergic interneurons and glutamate (NMDA) receptors that are involved in synaptic LTP and LTD^44–46^. By analyzing the learning progress, we could therefore disentangle whether a possible effect on memory retention was primarily driven by learning deficits already measurable during conditioning, or consolidation deficits only measurable during retention. However, our primary outcome measure was threat memory retention - which collapses learning and consolidation deficits - since cTBS may affect both processes and because the ultimate treatment outcome of attenuated aversive memories may be clinically most relevant.

## Results

### Threat memory retention: startle eye-blink response

On day 1 of the experiment, participants underwent cTBS immediately followed by discriminative threat conditioning with 50% reinforcement. On day 2, participants were assessed for threat memory retention (Fig. 1) in extinction (i.e., no reinforcement), immediately followed by a relearning session to assess memory savings from the conditioning session of the previous day. The primary focus here was to assess the cTBS effect on memory consolidation as assessed in the retention test. Consistent with a cTBS-induced reduction of threat memory retention, we found a significant group x CS type interaction (Table 1; Fig. 2), with overall smaller CS+/CS− difference in the experimental group compared to the control group. There was no evidence that this effect depended on CS complexity (Table 1). Post-hoc t-tests suggested a difference in fear-potentiated startle elicited by CS+ versus CS- (collapsing across CS complexity) in the control group, *t*(50) = 3.76, *p* = 0.0002, Cohen’s *d* = 0.53, but not in the experimental group, *t*(50) = 0.25, *p* = 0.60, *d* = 0.04.

**Figure 2.**
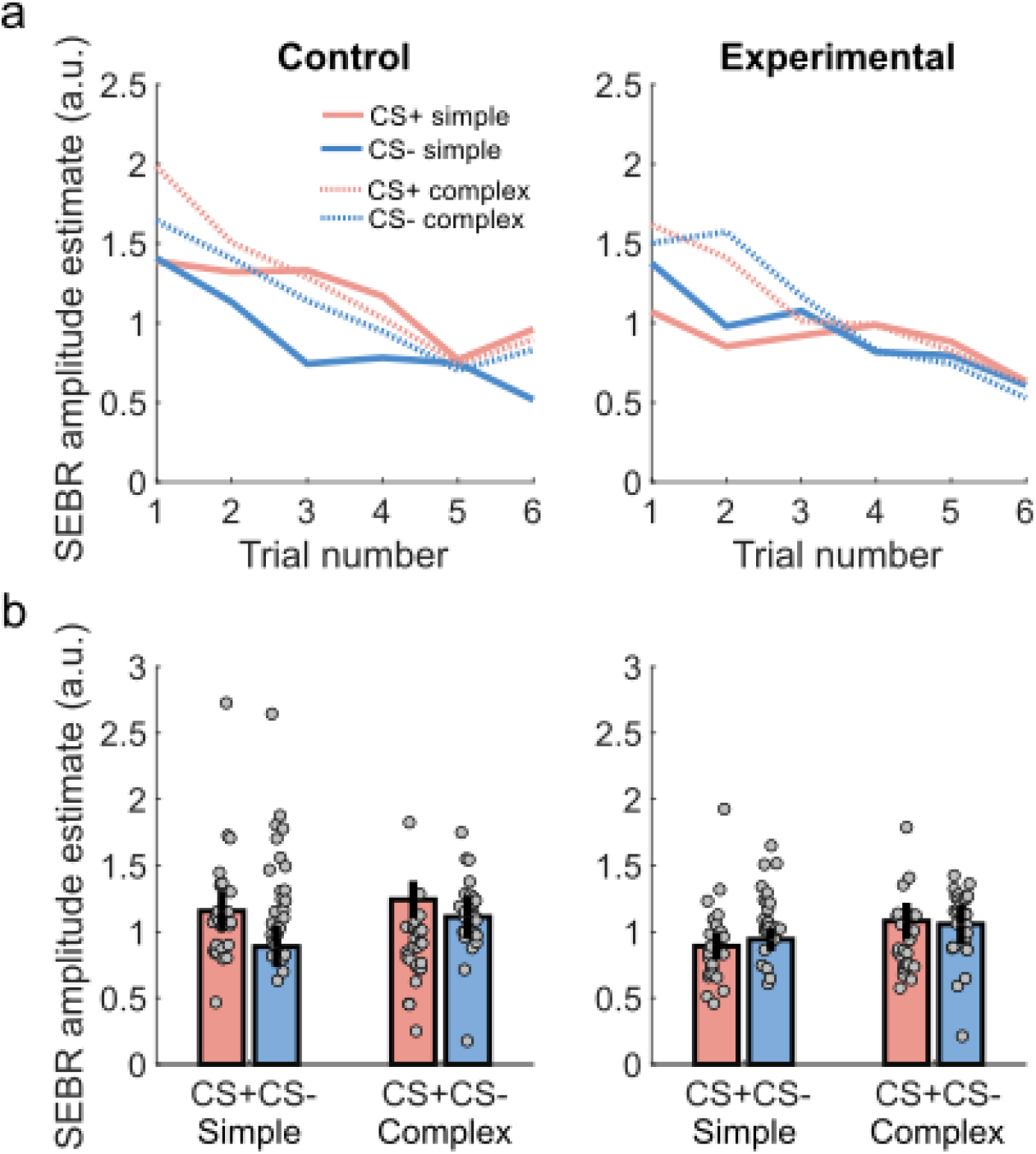
Startle eye-blink responses (SEBR) during threat memory retention test on test day 2. **a**. Trial-by-trial responses for each condition show the time course of the responses, with little differentiation of CS+ and CS– for the experimental group and overall quick decline in amplitude across trials (habituation). **b**. The CS+/CS- difference was smaller for experimental than for control group across simple and complex CS, indicating that the fear-potentiated startle was inhibited by continuous theta-burst TMS as measured in threat memory retention test the next day. Error bars are 95% within-subject standard errors of the mean reflecting paired, one-tailed CS+ > CS− comparison^47^; a. u. are arbitrary units.

**Table 1.** Statistical test results for condition-wise startle eye-blink responses during threat memory retention test on test day 2.

| Repeated-measures ANOVA | df1 | df2 | *F* | *p* | $\eta^2$ |
| --- | --- | --- | --- | --- | --- |
| Group | 1 | 50 | 7.71 | 0.008 | 0.024 |
| CS type | 1 | 50 | 6.39 | 0.015 | 0.020 |
| CS complexity | 1 | 50 | 5.48 | 0.023 | 0.049 |
| Group x CS type | 1 | 50 | 7.71 | 0.008 | 0.024 |
| Group x CS complexity | 1 | 50 | 0.00 | 0.99 | < 0.001 |
| CS type x CS complexity | 1 | 50 | 0.30 | 0.59 | < 0.001 |
| Group x CS type x CS complexity | 1 | 50 | 2.56 | 0.12 | 0.006 |

df = Degrees of Freedom, $\eta^2$ = Eta squared (explained variance).

This primary analysis was based on all trials of the retention test and may thus potentially reflect group differences in extinction learning during the retention test, due to repeated presentations of the CS without the US. However, detailed analyses showed that the group difference in CS+/CS- discrimination was already present during the first 6 trials of the retention test, *t*(49.71) = 2.16, *p* = 0.036, *d* = 0.598. Moreover, the size of the group difference in CS+/CS-discrimination did not differ between the first vs. the last half of the retention test, *t*(47.17) = 1.58, *p* = 0.12, *d* = 0.435.

Our primary analysis did not account for the distribution of CS+ and CS- across the random trial sequence, which by chance may have been different between the groups. To exclude such possible influences of this sequence randomness on habituation to the startle probe, we replicated the group x CS type interaction in a linear mixed effects model that took into account the linear effect of trial across conditions, and thus, potential habituation across time (interaction: *F*(1,1182) = 5.49, *p* = 0.019; Supplementary Table 2). Moreover, to ensure that our main result was not driven by outlier data points, we reanalyzed the data without two control group participants who had standardized startle amplitude larger than 2 (arbitrary) units for either simple or complex CS+ (Fig. 2b). The group x CS type interaction was less strong but remained statistically significant without these two participants, *F*(1,48) = 5.65, *p* = 0.021, *η*^2^ = 0.015.

### Threat conditioning on the day before retention test: skin conductance and pupil size responses

Our finding of reduced memory retention after cTBS could be due to impairments in synaptic consolidation after the threat conditioning protocol, or due to impairments in neural processing and synaptic transmission during the acquisition of conditioning. To assess the latter possibility, we analyzed skin conductance and pupil size responses to the CS during conditioning on test day 1. In line with our previous study that established the behavioral protocol for somatosensory threat conditioning^41^, participants learned to discriminate CS+ and CS- as evidenced by skin conductance responses (main effect of CS type; Table 2, Fig. 3a,c) and pupil size responses (Table 2, Fig. 3b,d). There was no clear evidence that learning differed between control or experimental cTBS (Group x CS type interaction) as reflected in SCR or PSR. Visual inspection of the SCR data may even suggest somewhat better, rather than worse, learning in the experimental group than in the control group (Fig. 3c).

**Figure 3.**
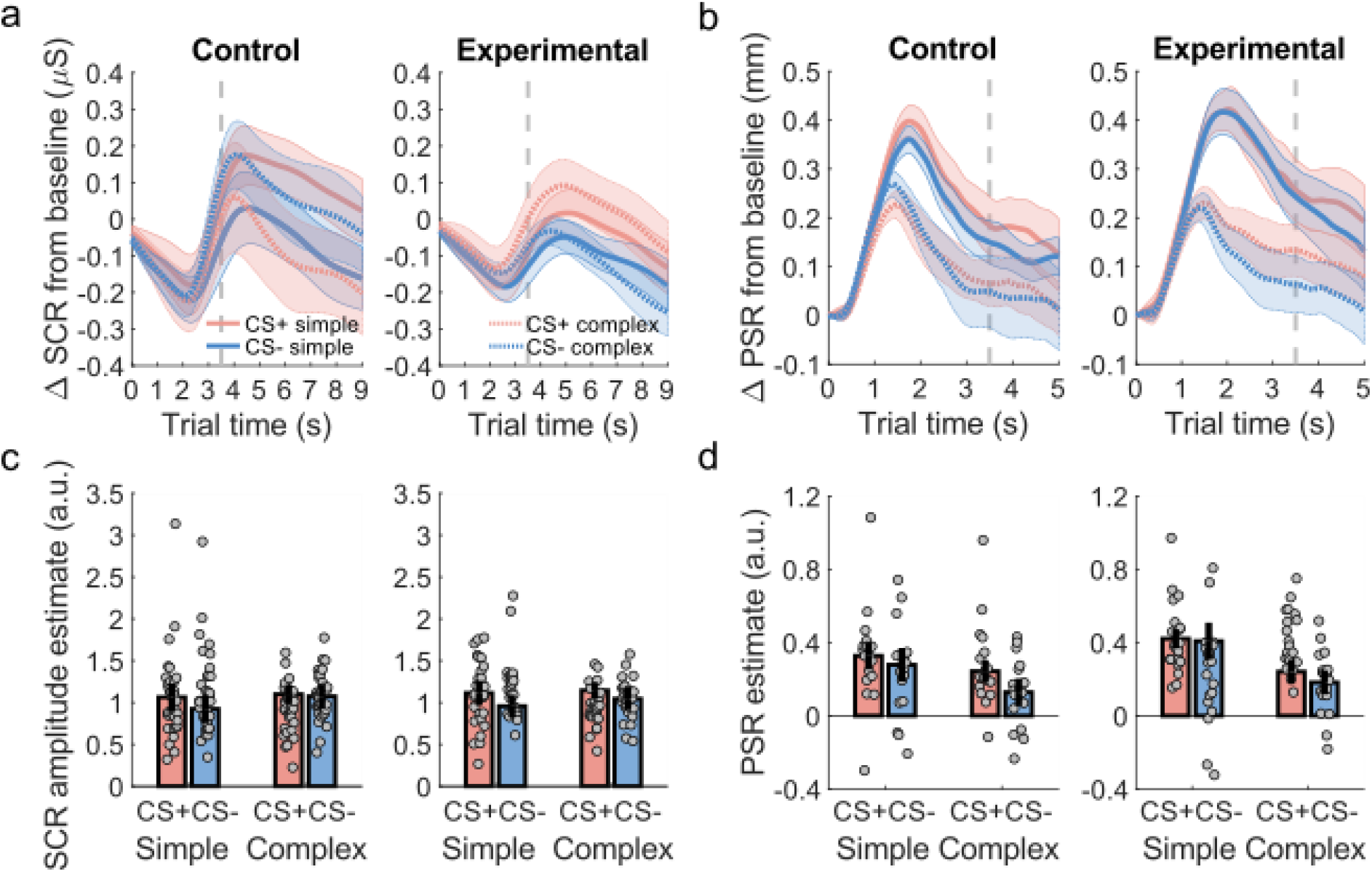
Skin conductance responses (SCR) and pupil size responses (PSR) during threat conditioning on test day 1. **a-b**. Averaged trial time courses of skin conductance and pupil size changes from baseline for the different stimuli, separately for control and experimental groups. Shaded areas represent within-subject standard errors of the mean; dashed grey line marks US onset (only no-US trials were included); *μS* are micro-Siemens. **b-c**. SCR amplitude and PSR estimates showed that the participants learned the CS+/CS- difference overall (main effect of CS type for both modalities, Table 2), evidencing threat conditioning. There was no evidence for a difference in learning between the control and experimental groups (interaction of group and CS type). Error bars show 95% within-subject standard errors of the mean reflecting paired, one-tailed CS+ > CS− comparison^47^; a. u. are arbitrary units.

**Table 2.**
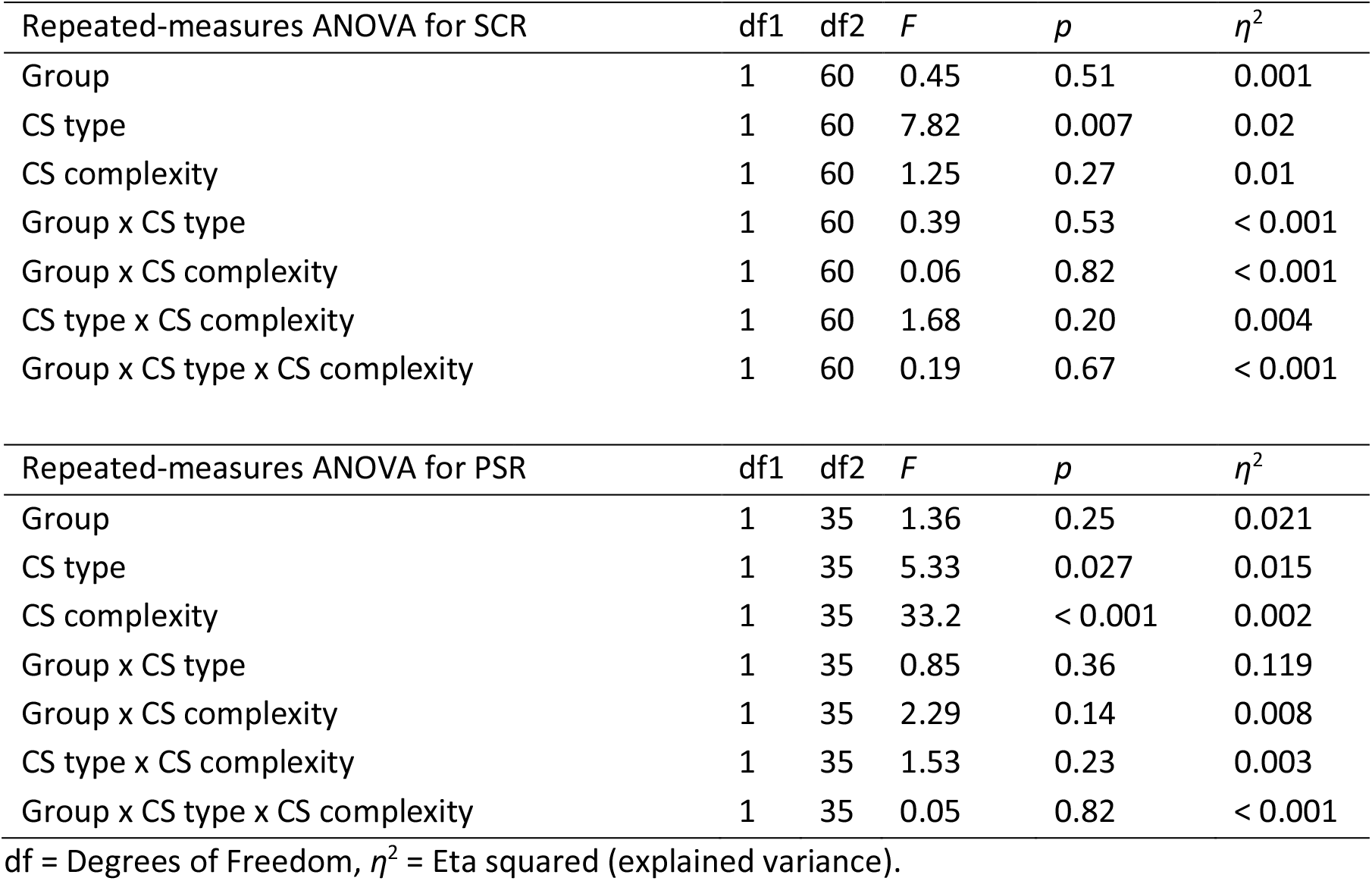
Statistical test results for condition-wise skin conductance and pupil size responses during threat learning on test day 1.

### Threat relearning after retention test: skin conductance and pupil size responses

To assess memory savings carried over from the conditioning session of the previous day, we conducted a second learning session directly after the threat memory retention test. Pupil size responses discriminated between threat-predictive CS+ and safe CS-, with no difference between experimental and control cTBS groups (Supplementary Fig. 1, Supplementary Table 5). No significant differences in CS+/CS- responses were found in condition-wise skin conductance responses (Supplementary Fig. 1, Supplementary Table 5) while a small difference was found in trial-wise responses, *F*(1, 3886) = 5.83, *p* = 0.02 (but no group x CS type interaction; Supplementary Table 6).

### CS-US contingency learning

We assessed declarative memory for the CS-US contingency using a continuous rating scale at the end of each test day. Participants distinguished CS+/CS-, *F*(1,50) = 18.1, *p* < 0.001, *η*^2^ = 0.121 (Supplementary Fig. 2a) after day 1 as well as on day 2 after relearning, *F*(1,50) = 83.3, *p* < 0.001, *η*^2^ = 0.356 (Supplementary Fig. 2b). There was no difference between ratings for simple and complex stimuli on either test day, or between cTBS groups on test day 1 (all *p* > 0.30). The CS type x CS complexity and group x CS type x CS complexity interactions after day 1 were not significant, *F*(1,50) = 3.49, *p* = 0.068, *η*^2^ = 0.001, and *F*(1,50) = 3.58, *p* = 0.064, *η*^2^ = 0.008, respectively. After test day 2, control cTBS participants discriminated CS+ and CS- better than experimental cTBS participants (group x CS type interaction: *F*(1,50) = 4.48, *p* = 0.039, *η^2^* = 0.019; Supplementary Fig. 2b). This difference was due to higher CS- ratings in the experimental group than in the control group, *t*(41.90) = −2.04, *p* = 0.048, Cohen’s *d* = 0.569. CS+/CS-discrimination in the control group increased from day 1 to day 2, *t*(26) = 2.39, *p* = 0.025, *d* = 0.46, whereas the increase in the experimental group was not statistically significant, *t*(24) = 1.80, *p* = 0.084, *d* = 0.361 (corrected α = 0.025). However, there was no difference between the groups in their increase in CS+/CS- discrimination from day 1 to day 2, *t*(46.82) = 0.86, *p* = 0.39, *d* = 0.237.

### Accuracy and reaction time of CS identification

As a control measure for potential cTBS-induced synaptic transmission impairments, which could lead to reduced stimulus perception, we analyzed accuracy and reaction times in CS identification. We found no evidence for reduced accuracy or increased reaction times in the experimental group in comparison with the control group. Accuracy was high in all sessions with 94.2% (± 2.7 percentage point SD) correct responses during conditioning, 96.9% (± 2.6 p.p.) during retention, and 95.8% (± 2.7 p.p.) during relearning. Accuracy was similar across groups and between CS+/CS- in all sessions (all *p* > 0.15). Participants were on average 4 percentage points more accurate in recognizing complex than simple CS during retention, *F*(1,50) = 7.43, *p* = 0.009, *η*^2^ = 0.03 (Supplementary Fig. 3a). There were no differences in reaction times across groups or CS type (all p > 0.50, except for CS type during relearning *F*(1,50) = 3.85, *p* = 0.056, *η*^2^ = 0.001). However, participants responded on average 620 ms faster to complex CS (mean RT 1670 ± 20 ms SD) than to simple CS (1050 ± 70 ms) during conditioning (*F*(1,50) = 228.9, *p* < 0.001, *η*^2^ = 0.448), 640 ms during retention (*F*(1,50) = 175.6, *p* < 0.001, *η*^2^ = 0.395; Supplementary Fig. 3b), and 610 ms during relearning (*F*(1,50) = 326.6, *p* < 0.001, *η*^2^ = 0.465). Importantly, this cannot be explained by physical properties or timing of the stimuli, as simple stimuli were immediately distinguishable from the first pulse, and therefore attentional or other psychological factors seem the most reasonable explanation for the difference (e.g., complex stimuli engaged more attention than simple stimuli). Average reaction times were similar to the conditioning session during both retention (simple CS 1680 ± 20 ms, complex CS 1040 ± 60 ms) and relearning (simple CS 1610 ± 70 ms, complex CS 1000 ± 40 ms).

## Discussion

Growing evidence from rodent studies suggests that sensory cortices are important for processing complex stimuli during auditory threat conditioning, but their role for simple conditioned stimuli (CS), and for human threat conditioning in general, remains unclear. We addressed this question in a transcranial magnetic stimulation study with the goal of inducing temporary inhibition of neural processing in the sensory cortex. We applied continuous theta-burst stimulation (cTBS) in healthy human participants over the primary somatosensory cortex, either ipsilateral (control) or contralateral (experimental) to somatosensory stimuli, immediately prior to a threat conditioning protocol where the somatosensory stimuli were paired with painful shocks (CS+) or not (CS-). We found that after overnight consolidation, differential fear-potentiated startle to the CS+/CS- was smaller in the experimental (contralateral cTBS) group compared with the control (ipsilateral cTBS) group. Detailed analyses confirmed the robustness of this finding and showed that it was not due to group differences in extinction learning or in the random trial sequence. We found no evidence for an inhibitory cTBS effect on learning during the initial learning session, indicated by a lack of group differences in how skin conductance and pupil size responses discriminated between CS+ and CS-. Moreover, there was no evidence that contralateral cTBS impaired CS identification, excluding the possibility that our results reflect a cTBS-induced deficit in immediate somatosensory processing unrelated to the threat conditioning. Taken together, the observed effect on threat memory retention suggests that the cTBS did not interfere with basic sensory processing and activity-dependent short-term plasticity, but rather with synaptic structural reconfiguration required for memory consolidation^48^.

A few previous studies have used TMS to investigate the role of cortical areas in threat conditioning. These studies have mainly focused on TMS effects during memory retrieval and extinction, and none have investigated the role of sensory cortices. One study found that 20 min of inhibitory 1-Hz repetitive TMS targeting right posterior parietal cortex after threat conditioning increased reaction times in a subsequent visual search task when a CS+ was presented as a distractor stimulus, potentially reflecting impaired disengagement of attention due to the TMS intervention^49^. Recently, it was shown that 15 min of 1-Hz repetitive TMS over dorsolateral prefrontal cortex administered after visual threat memory retrieval was successful in reducing physiological threat responses during the so-called reconsolidation period, as well as preventing return of threat responding after reinstatement, while leaving declarative CS-US learning intact^50^. The authors speculate that this effect may be due to the role of dorsolateral prefrontal cortex in memory retrieval, for example through working memory processing of retrieved information, and potentially due to long-range connections from the prefrontal cortex to the amygdala^50^. Therefore, there may be several putative time windows and neural pathways through which to influence threat memories with TMS. A few other studies have targeted extinction learning mainly through ventromedial prefrontal cortex stimulation^51–54^. Beyond TMS studies, there is very little data from clinical lesion samples to address the role of sensory cortices in threat conditioning. One lesion case study suggested that visual cortex is not required for non-discriminative visual conditioning in humans^55^, in line with non-discriminative auditory conditioning in rodents^12,14,16–18^.

We selected somatosensory threat conditioning as a model system for methodological reasons. We note that somatosensation is crucial for adaptive functioning and is therefore also an important sensory modality to investigate, next to the more commonly investigated auditory and visual modalities. Clinically, somatosensation is affected in post-traumatic stress disorder^56^, a psychiatric disorder that has been proposed to develop partly due to maladaptive conditioning mechanisms such as threat overgeneralization and impaired safety learning^1,2^. Somatosensory conditioning has been behaviorally demonstrated across species. In rodents, somatosensory threat conditioning with whisker touch as CS is suggested to induce increased neural response strength, sparse coding^57^, dendritic spine plasticity^58^, and inhibitory postsynaptic potentials^59^ in the primary somatosensory barrel cortex, somewhat similar to post-consolidation changes in the auditory cortex^60^. In humans, somatosensory threat-conditioned CS slightly increase the painfulness of the US^39^. Moreover, specificity of threat responses is affected by somatosensory precision of the body part, with differential conditioning observed for CS applied on the hand but not for CS on the back^40^. Our study adds to the scarce literature on somatosensory threat conditioning in humans. In line with our previous work^41^, we have shown that somatosensory threat learning is possible from simple and complex patterned electric pulses on fingers, but we now additionally demonstrate that the associative memory is retained at least overnight.

We found that primary somatosensory cortex (S1) is required for consolidation of associative somatosensory threat memories in humans, in line with previous findings in rodent auditory threat conditioning to complex cues. It has been suggested that primary sensory cortices are involved in stimulus identification and are specifically needed for processing complex stimuli, because they necessitate the binding together of different stimulus elements into a unitary representation, for example frequency sweeps and tone pips^22,61^. However, we did not find clear evidence that the role of S1 in threat conditioning and threat-memory retention differed between simple and complex somatosensory stimuli (here differentiated by the temporal pattern of the stimulus across one or two fingers). On the other hand, our findings are in line with human neuroimaging studies that found equal CS+/CS- neural pattern discriminability for simple and complex CS in the auditory cortex^34,35^ and that were able to cross-decode simple CS threat predictions from complex threat predictions and vice versa^34^. However, it is possible that experiments with larger sample sizes may yet detect such complexity effects^62,63^.

Secondary sensory cortices have been shown to be involved in learned threat predictions of auditory, visual and olfactory modalities in rodents^12^. The effect of stimulus complexity has not been investigated across modalities, and to our knowledge, no comparison including somatosensory threat conditioning exists, nor has the role of primary sensory cortices been compared across modalities. TMS is known to influence wide functional networks in the human brain, with potential effects on even distant brain regions through long-range connections^43,64^. This has also been shown for cTBS of the primary somatosensory cortex at rest, which led to reductions in functional connectivity of S1 with a variety of distant brain regions, including regions important for threat conditioning such as the amygdala, striatum and anterior cingulate cortex^65^. Therefore, without concurrent fMRI measurement, we cannot confirm or exclude possible downstream influences of our cTBS protocol on secondary somatosensory cortices or other areas, although the stimulation is unlikely to have directly reached the ipsilateral secondary somatosensory cortex.

The neural effects of theta-burst TMS are multifaceted^42^. They may reflect synaptic long-term potentiation and depression at excitatory glutamatergic^44,66^ and inhibitory GABAergic neurons^46,67^, neurotrophic factors^68,69^, and neurogenesis^70^. It has been proposed that the long-term depression (LTD)-like effect of continuous theta-burst TMS used here is based on an initial facilitatory long-term potentiation (LTP)-like effect, as seen with intermittent theta-burst TMS, which reverses and becomes inhibitory with continuing stimulation^71^. This build-up of the inhibitory effect may be determined by post-synaptic calcium levels that act as a trigger for synaptic plasticity for both facilitation and inhibition, whereby the sum of LTP and LTD together determines the final direction of the TMS after-effect (facilitatory or inhibitory)^71^. The evidence comes from in vitro experiments on rat hippocampal slices^72^ and computational models proposing that extracellular stimulation up to 5 Hz induces longterm depression with moderate levels of post-synaptic calcium, controlled by NMDA receptor activity^73^. However, at this moment it is not yet known whether these mechanisms truly underlie the observed inhibitory effect of continuous theta-burst TMS observed in humans. The best evidence thus far comes from a study where an NMDA-receptor antagonist blocked both the LTP-like excitatory and LTD-like inhibitory effects of theta-burst TMS in six healthy human participants^44^. Finally, the efficacy of repetitive TMS is crucially influenced by baseline cortical excitability as well as individual and contextual factors^42,45,74^. While there are thus several potential low-level physiological mechanisms that may have led to the effects observed in our study, our results suggest that cTBS can selectively influence memory retention in S1 without strong effects on immediate sensory processing and activity-dependent short-term learning.

Taken together, we found that inhibitory continuous theta-burst TMS over the primary somatosensory cortex led to reduced threat memory retention after overnight consolidation, with no clear influence of conditioned stimulus complexity. This finding extends the rodent and human literature on the function of primary sensory cortices in threat learning and memory, by suggesting that primary sensory cortices are involved in stimulus representation for threat conditioning and play a role in reconsolidation of threat memories. Further studies may investigate the neurophysiological mechanisms underlying these effects with concurrent neuroimaging and TMS, examine whether continuous theta-burst TMS on primary somatosensory cortex could be used as a method to modify labile reactivated threat memories through reconsolidation update mechanism^75^, and attempt to establish whether intervening with (re)consolidation of threat memories may have clinical relevance for patients suffering from fear and anxiety disorders.

## Methods

### Participants

Sixty-eight participants (34 women and 34 men; mean age 23.7±4.2 SD) recruited from the general and student population completed the cTBS and threat conditioning session on test day 1 according to protocol. After exclusions due to data collection and quality issues (detailed in the respective sections below), the final sample for skin conductance analyses was 62 participants (28 experimental contralateral cTBS group, 34 ipsilateral control cTBS group). For pupil analyses, the final sample was 34 participants (18 experimental, 16 control participants). Sixty-two participants completed the threat memory retention session on test day 2 according to protocol. Fifty-two participants (26 women and 26 men; mean age 23.8 ± 3.9 SD) were included in our main analyses of startle-eye blink responses in the threat memory retention test. Twenty-five of these participants (12 women, 13 men) had been randomly assigned to the experimental (contralateral cTBS) group and 27 (14 women, 13 men) to the control (ipsilateral cTBS) group. Sixty-five participants completed the relearning session on test day 2 according to protocol. The final sample for skin conductance analyses comprised 56 participants (27 experimental, 29 control) 42 participants for pupil size analyses (22 experimental, 20 control). Twenty additional participants took part in an initial fMRI session but were excluded from TMS due to inability to localize S1.

Details of the exclusions are given in the *Experimental Procedure* section. All recruited participants stated no history of neurological and psychiatric disorders or contraindications for TMS and MRI and gave written informed consent. The study protocol, including the form of taking consent, was in accordance with the Declaration of Helsinki and approved by the governmental research ethics committee (Kantonale Ethikkommission Zürich, KEH-ZH 2013-0383).

### Experimental procedure

#### fMRI localizer for S1

Participants took part in an fMRI experiment to localize the S1 representation of index and middle fingers of both hands. Somatosensory stimulation to each finger was delivered with a constant current stimulator (Digitimer DS7A, Digitimer, Welwyn Garden City, UK) through a pin-cathode/ring-anode electrode. Each stimulation consisted of a 4-s train of 20 square electric pulses of 200 μs (in 4 bursts of 5 pulses, with ~10 ms breaks between pulses and 150 ms breaks between bursts), on one finger at a time. Intensity of the stimulation was individually calibrated for each participant and each finger before the experiment by incrementally increasing the stimulator current until the participant indicated that the intensity was clearly perceptible but not yet unpleasant. The localizer experiment contained three fMRI runs and the somatosensory stimulation intensity was checked between runs and re-calibrated if necessary. Participants were instructed to lie still in the scanner and focus on the sensation in their fingers. Somatosensory stimulation was delivered in the first two runs repeatedly in order from left middle finger through index fingers to right middle finger with a 6-9 s interstimulus interval (ITI) on 26 trials per run. In the third run, stimulation was delivered in pseudorandom order (max. 3 consecutive trials of one finger) with 2-4 s ITI on 43 trials. Each run lasted 300 seconds and the experiment had in total 95 trials. The experiment was programmed with Psychophysics Toobox^76^ in MATLAB2014b. Functional images were acquired with a Philips Achieva 3 T MRI scanner with 8-channel sensitivity-encoded (SENSE) head coil. T_2_^*^-weighted echo-planar images were acquired with the following parameters: 3 x 3 x 3 mm voxel size, 33 ascending axial slices, 0.5 mm gap, volume TR 2000 ms, 150 volumes per run, TE 30 ms, flip angle 90°, FOV 240 x 240 x 115 mm, and acquisition matrix 80 x 78, 5 dummy scans. A high-resolution structural scan was obtained after the three functional runs (MP-RAGE; 1 x 1 x 1 mm voxel size, 181 sagittal slices).

We used SPM12 (Wellcome Trust Centre for Neuroimaging, London) and MATLAB2016a to preprocess and analyze fMRI data. Preprocessing of the functional images included realignment, slice-time correction^77^, co-registration with the T_1_-weighted structural images, normalization to the Montreal Neurological Institute (MNI) space based on unified segmentation^78^, and smoothing with an 8 mm FWHM Gaussian kernel. Each trial of somatosensory stimulation was modelled as a stick function of zero duration at trial onset, which led to the most stable effects (vs. a boxcar model with duration of 4 seconds). Contrasts were defined for ‘left > right hand’ and ‘right > left hand’ BOLD activity. For each participant and each hemisphere, the stimulation site ROI for cTBS was defined as a 10-mm diameter sphere around the peak coordinates within an anatomical S1 mask (Fig. 4). The S1 mask was created as an intersection of anatomical postcentral gyrus mask from Harvard-Oxford cortical and subcortical structural atlases (fsl.fmrib.ox.ac.uk/fsl/fslwiki/Atlases), a 15 mm radius sphere around MNI coordinates from a finger somatotopy study^79^, and grey matter mask from the segmentation procedure at 0.5 threshold. We included participants with activity of T ≥ 1.28 (corresponding to p < 0.01 uncorrected) for both ‘left > right’ and ‘right > left’ contrasts. We excluded those participants whose peak BOLD activity was located very inferior toward the lateral sulcus and/or deep within a sulcus, as there was a risk that these locations correspond to S2 rather than S1, and because it is difficult to reliably target such sites with our TMS approach. In total, these criteria excluded 20 fMRI participants from the TMS session. Following this analysis, each participant was randomly assigned to experimental or control group by a researcher outside the study team.

**Figure 4.**
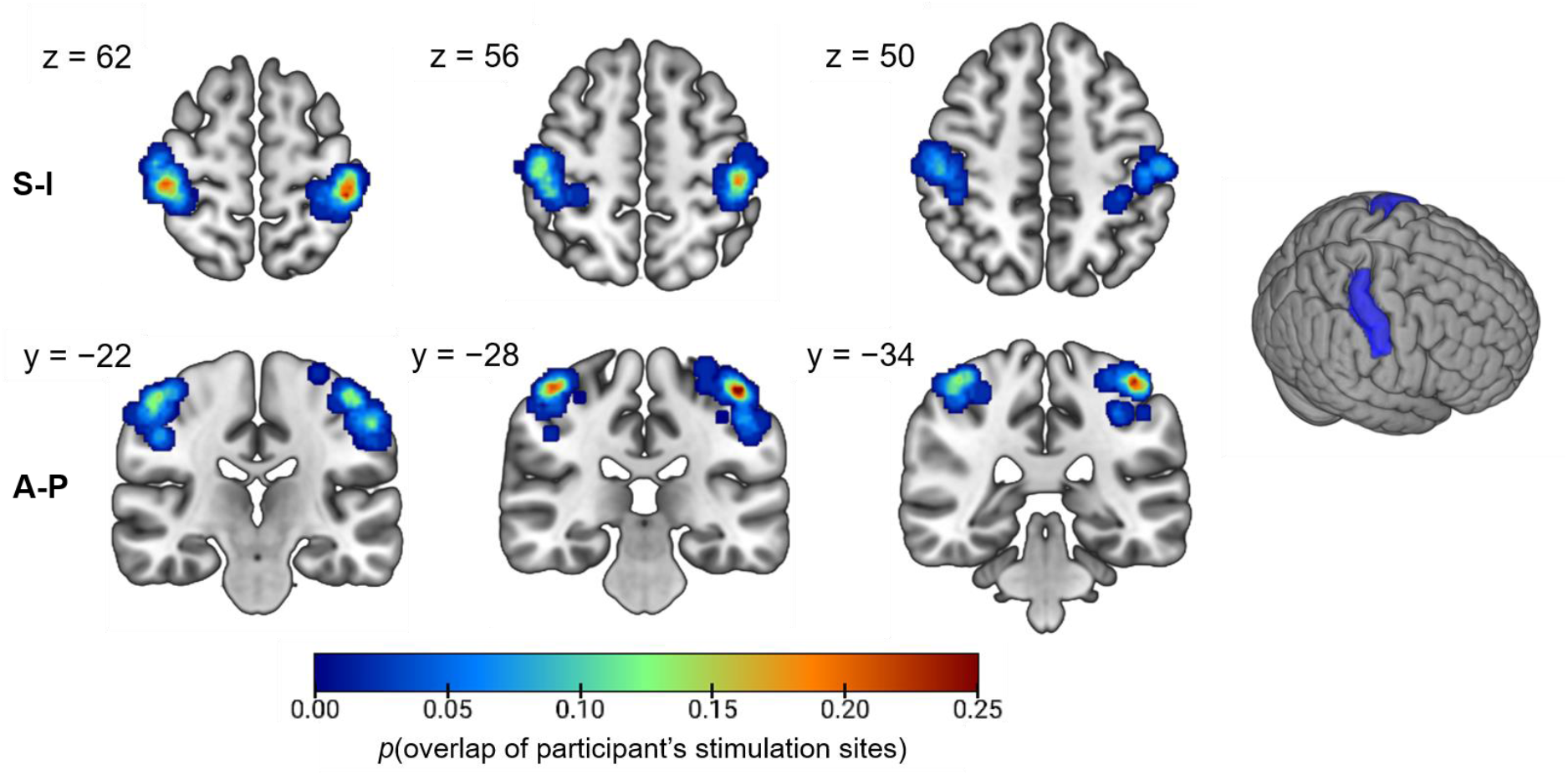
Probability map of S1 stimulation site ROIs across all participants for both hemispheres overlaid on T1 in MNI152 space. On the right side is the S1 mask in the MNI space. A-P: Anterior-Posterior, S-I: Superior-Inferior. Color indicates the probability (0-1) of an overlap of the participants’ stimulation site ROIs at each coordinate.

#### Continuous theta-burst TMS protocol

After the fMRI localizer, participants were invited to the laboratory on two consecutive days (Fig. 1). On the first day, continuous theta-burst repetitive TMS was administered before the experimental task. We used an inhibitory protocol that has been shown to decrease cortical excitability for up to an hour, as measured with motor-evoked potentials^37^. The S1 ROI coordinates in MNI space from the fMRI localizer analysis were inverse transformed to each participant’s native anatomical T1 space in SPM12 and overlaid on a constructed 3D model of the participant’s head and brain using the BrainSight interface (Rogue Resolutions Ltd., Cardiff, UK). For the experimental group, cTBS was applied to the left hemisphere ROI, and for the control group to the right hemisphere ROI. The precise individual stimulation site was defined as the approximate center of the S1 spherical ROI while attempting to also optimize the location so that 1) the TMS coil was angled perpendicular to the crown of the postcentral gyrus, 2) the coil surface was parallel to both the brain and the skull at the stimulation site, 3) the coil handle pointed backward. The cTBS site was targeted with neuronavigation (BrainSight and Polaris NorthernDigital, Waterloo, Canada infrared tracking system). For relating the anatomical image of the participant in BrainSight to the dimensions of the participant’s head in real space, landmarks were set on bridge of the nose, tip of the nose, and bilateral intratrageal notches at the ears. The position of the participant’s head and the coil were tracked with the Polaris system for accurate pulse delivery to the stimulation site.

cTBS intensity for each participant was defined via a standard motor threshold procedure with single-pulse TMS on the index finger motor cortex. The passive motor threshold was defined as the intensity that elicited motor evoked potentials of > 200 μV in amplitude in at least 5 out of 10 pulses as measured by electromyography of the first dorsal interosseous muscle between the index finger and the thumb. For obtaining the active motor threshold, the participant was asked to exert constant pressure between the index finger and the thumb (20% of maximum force) while the procedure described above was repeated. Intensity of the cTBS on the S1 ROI was set to 80% of the active motor threshold, scaled by the difference between the distance of the motor cortex and the distance of the S1 ROI to the skull^80^. The average stimulation intensity across participants (N = 68 with TMS completed according to protocol) was 41.3% (SD 6.7, range 25-51%) of maximum stimulator output.

The cTBS protocol consisted of 600 pulses administered continuously for 40 seconds in bursts of three pulses at 50 Hz (every 20 ms) at 5 Hz intervals (every 200 ms)^37^. The stimulation was applied with a Magstim super-rapid stimulator (Magstim, Whitland, Dyfed, UK) and a figure-of-eight coil, supported by a frame over the stimulation site and held steady by hand. Immediately after cTBS, participants underwent 20 minutes of classical threat conditioning, which is well within the estimated duration of the cTBS effect^37^.

#### Conditioned and unconditioned stimuli of the threat conditioning protocol

There were four different CSs in the experiment: 1) simple CS+ paired with an electric shock US in 50% of trials, 2) simple CS- never paired with the US, 3) complex CS+ paired with an electric shock US in 50% of trials, 4) complex CS- never paired with the shock. Simple and complex CS trials were presented in separate, alternating conditioning blocks in all conditioning and retention sessions. The intensity for the CS was set for each finger separately and to a subjectively clear but not unpleasant level by incremental increases of the current. CS were 4-s electric pulses to the middle and index fingers of the left hand delivered with Digitimer DS7A stimulator through a pin-cathode/ring-anode electrode. All CS consisted of square electric pulse trains, with simple stimuli delivered as 20 pulses of 200 μs (in 4 bursts of 5 pulses, with ~10 ms breaks between pulses and 150 ms breaks between bursts) to one finger only. Complex stimuli were 24 pulses of 200 μs in 3 bursts of 5 pulses per finger, with ~10 ms breaks between pulses and 450 ms breaks between burst sets, alternating from one finger to another. The complex stimuli always started with the same finger within a participant (randomized across participants), so that recognition of the stimulus was only possible after the first 3 bursts. One complex stimulus would contain 3 bursts on one finger first, then a break of 450 ms, and then continue with 3+3 alternating pulses on the two fingers and end with 3 pulses on the other finger, whereas the other complex stimulus would alternate 3+3 bursts on the two fingers throughout (Fig. 1).

Aversive US was an electric shock consisting of 50 square pulses of 200 μs for 500 ms, with ~10 ms between each pulse, delivered with Digitimer DS7A stimulator to the top of the foot, approximately on the intermediate cuneiform bone. The US was delivered ipsilateral to the cTBS site in order to exclude possible cTBS effects on sensory processing of the US. Intensity of the US was individually calibrated for each participant. First, a clearly unpleasant but not painful intensity was determined with an ascending staircase procedure. After that, participants gave subjective ratings (0 = felt nothing to 100 = very unpleasant) for 14 random intensities. The 90% intensity based on these ratings was chosen as the US intensity for the experiment (N = 68, 6.1 ± 5.7 mA, range 1.5–41 mA).

#### Day 1: training, cTBS and threat conditioning

Participants trained the experimental task without reinforcement before cTBS. They were exposed to each of the 4 somatosensory conditioned stimuli (CS) three times in random order (i.e., 12 trials) and were asked to indicate which CS pattern they perceived by pressing a left or right button during 3 s from CS onset. This training block was repeated until the participant achieved 75% accuracy (i.e., 4/6) both for simple and for complex stimuli, and served also as habituation to the CS alone.

After cTBS, the 20-minute threat conditioning protocol followed, consisting of 8 blocks of 12 trials each, for a total of 96 trials. The intertrial interval for each trial was drawn randomly from a uniform distribution ranging from 7 to 11 seconds in steps of one second. The order of simple and complex blocks (simple first or complex first) was randomized. Due to a technical error in the procedure to balance the stimulus complexity order across participants, about 14% of participants received simple stimuli first and 86% of participants received complex stimuli first. However, the number of participants that received simple first or complex first were balanced between experimental and control groups. Out of the participants that were included for the retention session startle-eye blink analyses, 45 participants received complex stimuli first and 7 simple stimuli first, and of those, 35 participants received middle finger stimulation as simple CS+ and leftward wave as complex CS+, and 17 participants had the index finger for simple CS+ and rightward wave as complex CS+. 37 participants had their complex stimuli starting from the middle finger and 15 participants from the index finger.

The experiment was run in Psychophysics Toolbox^76^ in MATLAB2013b. The visual part of the experiment was shown on an 18.5” screen (1280 x 1024 pixels resolution) at approximately 90 cm distance from the participants’ eyes. The participants were instructed to fixate their gaze on a white cross on grey background during the trials. There was no other visual element on the screen during the trials.

After the conditioning session, seven participants were excluded from all analyses due to either a failure to complete the cTBS according to protocol (e.g., the participant and therefore the coil moved during the 40 seconds of stimulation, or the BrainSight program had a technical failure), or due to failure to deliver the US throughout the conditioning session (e.g., electrode detachment).

#### Day 2: threat memory retention and relearning

The next day, participants returned for a retention and relearning session. Retention was measured under extinction (i.e., no US) in 4 blocks of 6 trials each, i.e., 2 blocks of simple and 2 blocks of complex stimuli, for a total of 24 trials (12 CS+, 12 CS-). Before the retention session, the experiment was set up the same way as the previous day, including the US electrode on the foot. In order to retain expectation of US, participants were instructed that they may experience US during the session, including the possibility of only one US at the very end. During the retention session, participants wore headphones from which they heard a 50-ms, 100 dB burst of white noise on every trial at CS offset to elicit an acoustic startle reflex. Immediately before the retention test started, one habituation startle probe was presented alone. After the retention session, headphones were taken off and the participants underwent a re-learning session with the same structure and duration as the conditioning session on the previous day, including US delivery.

### Data recording and psychophysiological modelling

#### CS-US contingency ratings

Immediately after the conditioning and re-learning sessions, participants were presented with each CS alone, without US. Participants were asked to indicate what they thought was the probability of the US occurring together with this CS, from a choice of 0%, 25%, 50%, 75%, or 100%.

#### Skin conductance responses

We measured learning progress during conditioning and relearning by recording skin conductance from the first phalanx of the ring and little fingers of the left hand with plate electrodes. Skin conductance responses were not analyzed for retention on test day 2 due to large signal changes and artefacts caused by the startle probes and associated startle movements. The skin conductance signals were recorded with PowerLab 4/25T system (ADInstruments, Sydney, Australia), digitised (sampling frequency 1000 Hz) and saved in LabChart software (ADInstruments). Data files were converted into MATLAB format in LabChart and imported into PsychoPhysiological Modelling (PsPM)^81^ toolbox (version 4.1.1, bachlab.github.io/PsPM) in MATLAB2018a for further preprocessing and analysis. Data were visually inspected, and artefacts excluded (e.g., electrode movement or detachment). For analysis of the conditioning session, 3 participants were excluded due to recording failure (e.g., missing data or markers), and 3 were excluded due to data quality issues (e.g., continuous movement or detachment of the plate electrodes). For the relearning session, we excluded 8 and 4 participants, respectively, by the same criteria. Data were filtered with a first order bidirectional bandpass Butterworth filter with cutoff frequencies of 0.0159 and 5 Hz, and down-sampled to 10 Hz.

We used the PsPM toolbox for psychophysiological modelling of the skin conductance responses, involving inverting a nonlinear dynamic causal model (DCM)^82,83^ that estimated the amplitude of anticipatory (conditioned) and evoked (unconditioned) responses based on a canonical response function developed in earlier work^84–86^. For plotting of condition-wise averages of skin conductance responses over trial time, the signal was down-sampled to 50 Hz and baseline-corrected by subtracting the average signal from the 1 second time interval preceding trial onset^83^.

#### Pupil size responses

Pupil size and gaze direction were recorded during conditioning and re-learning with an EyeLink 1000 System at 500 Hz sampling frequency (SR Research, Ottawa, ON, Canada). The eye-tracker was placed at 80 cm distance from the subject’s eyes. To avoid recording issues associated with the electromyography electrodes placed directly underneath the eye (which were not removed for the relearning session after they were used to measure the startle eye-blink responses during the retention session, see next section), only the right eye was recorded. Pupil size responses were not analyzed for retention on test day 2 due to eye-blinks caused by the startle probe. Calibration of gaze direction was conducted on a 3-by-3-point grid in the EyeLink software. EyeLink data files were converted into .asc format and imported into PsPM toolbox (version 4.1.1 and 4.3.0) in MATLAB2018a for further preprocessing and analysis. Two participants were excluded from the conditioning session analyses due to pupil size and/or gaze direction data not recorded according to protocol (e.g., missing data or markers), and 6 were excluded from the relearning session analyses. During import into PsPM, blink and saccade periods detected by the EyeLink online parsing algorithm were excluded. Pupil size data points for which gaze direction deviated more than 7° visual angle from the center of the screen were excluded^41,87^. After these exclusions, more than 50% of pupil size data was missing during the 3.5 seconds of CS presentation before US onset for 32 participants in the conditioning session, and for 20 participants in the re-learning session. This is most likely because participants did not comply with the instruction to look at the (static) screen and keep their eyes open. Indeed, participants were often observed by experimenters to look away from the screen specifically during the somatosensory CS stimulation, as if to focus on the sensation. Therefore, only 34 participants that had at least 50% remaining data for each of the 8 blocks and were included in the pupil size analyses for the learning session and 42 for the re-learning session.

We used the PsPM toolbox to invert a general linear model based on a canonical pupil size response for threat conditioning and the measured data to arrive at estimates of condition-wise pupil size responses^41^. Missing data points (due to blinks, saccades, or loss of fixation) were interpolated for filtering and ignored during model inversion. Prior to model inversion, data were filtered with a bidirectional first order Butterworth low pass filter with 50 Hz cut off frequency and down-sampled to 100 Hz. For plotting of condition-wise averages of skin conductance responses over trial time, the signal was baseline-corrected by subtracting the first data point after trial onset^41^.

#### Startle eye-blink responses

The startle eye blink reflex was measured during the retention session using electromyography (EMG). The startle eye-blink was chosen as the main measure of interest since it allows the lowest effect size of all psychophysiological measures to detect at least 50% reduction in differential conditioned response at alpha level 0.05 for a one-tailed t-test (CS+ > CS-)^62,63^. However, startle-eye blink data was not collected from the acquisition (nor relearning) session due to the disruptive properties of acoustic startle probes on skin conductance and pupil size signals mentioned above, next to evidence that it slows down threat acquisition^88^. Two EMG electrodes filled with conductive gel were placed on the *orbicularis oculis* muscle under the left eye, approximately 1 cm apart^89^. The EMG and startle probes were recorded with a PowerLab 4/25T system (ADInstruments, Sydney, Australia), digitised (sampling frequency 1000 Hz) and saved in LabChart software (ADInstruments).

Data files were converted into MATLAB format in LabChart and imported into PsPM toolbox (version 4.3.0) in MATLAB2018a for further preprocessing and analysis. Trial markers were constructed from acoustic startle sound data channel with PsPM function pspm_find_sounds. EMG data pre-processing followed the optimized method reported in earlier work^90^. EMG data were filtered with a 4^th^ order Butterworth filter with 50 Hz and 470 Hz cutoff frequencies, and mains noise was removed with a 50 Hz notch filter. Then, the data were rectified and smoothed with a lowpass filter with 53.05 Hz cutoff (3 ms time constant). Ten participants were excluded based on visual inspection of overall trial-averaged EMG data. These participants had extremely noisy average startle eye-blink responses that were also often contaminated by CS stimulation artefacts.

Trial-wise startle eye-blink responses were estimated with a flexible-latency general linear model for each participant, using the PsPM toolbox^90^. The latency of the response was set to vary freely and the time window over which the model was evaluated was set at −0.02 and 0.13 s relative to the onset of the sound marker for each trial^90^.

### Statistical analysis

The estimated startle eye-blink, skin conductance and pupil size responses, and post-experiment contingency ratings, accuracies and reaction times for each participant were exported to and analyzed in R^91^ (version 4.0.2) run with RStudio^92^ (version 1.3.959).

As in previous work^62,93^, skin conductance responses from the conditioning session and startle-eye blink responses from the retention session were standardized within-subjects with respect to the average CS- response across conditions to account for different scaling. This was not done for pupil size responses due to smaller between-subjects variability in the individual pupil size response. The standardized trial-wise responses were averaged over trials within each condition for condition-wise analyses. For skin conductance responses and pupil size responses in both learning and re-learning sessions, only trials without US reinforcement were included in the analyses to avoid confounding of the conditioned responses with unconditioned responses. Trial-wise response estimates for skin conductance responses were averaged over trials. Pupil size responses were estimated on a condition-by-condition basis. Reaction times and accuracy percentages were analyzed for all sessions and CS-US contingency ratings from after conditioning and after re-learning. Trials with reaction times shorter than 200 ms and longer than 3500 ms were removed from analyses to exclude contamination of the results by either late responding from previous trial or responding after US onset at 3.5 s.

Psychophysiological response estimates, CS-US contingency ratings, reaction times and accuracy percentages were entered into three-way repeated measures ANOVAs of the form DV ~ Group * CS type * CS complexity + Error(Subject/(CS type * CS complexity), including all possible two- and three-way interactions. Post-hoc tests were conducted with emmeans R package^94^ and effect sizes for the post-hoc contrasts were approximated with ‘t_to_d’ function of effectsize package^95^. Further specific tests were conducted with Student’s paired t-tests for within-subject effects and with Welch’s two-sample t-tests for comparing groups with unequal sample sizes (including a correction for degrees of freedom). One-tailed p-values were used for a priori directional hypotheses for CS+ > CS- and control > experimental group; two-tailed tests were used for all other comparisons.

For startle eye blink responses, the random trial sequence and startle response habituation may inadvertently influence condition averages: for example, if a participant has more CS+ trials early in a block, then due to habituation, the CS+ average will be higher than the CS- average. To account for this possibility, we conducted a control analysis using a linear mixed effect model (LME; nlme package^96^) with the trial index across conditions as fixed factor. Thus, this model contained the within-subject factors Trial (linear effect of trial number across conditions), CS type (CS+ vs. CS-) and CS complexity (simple vs. complex), and between-subjects factor cTBS Group (experimental vs. control), as well as all possible interactions (up to four-way interactions). To formally compare possible models with different random effects, we computed log Bayes Factors (BF) with Bayesian Information Criterion approximation for frequentist linear regression models with the R package bayestestR^97,98^. The following random effects structures were estimable and compared to each other: 1|Subject as the null model, and (CS type * Trial)|Subject. The null model with 1|Subject as random effects was selected as the other model fit the data worse (log BF = −19.7). As a control analysis, trial-wise skin conductance responses were also analyzed with a linear mixed model (nlme package) similar to the one for startle eye-blink responses. For skin conductance responses, only the simplest possible random effects structure, 1|Subject, was estimable for both the conditioning and relearning sessions.

## Supporting information

Ojala et al. 2021 Supplementary Information

Figure 2 / Table 1 / Supplementary Table 1 Summary data 1

Figure 3 / Table 2 / Supplementary Table 3 Summary data

Supplementary Table 2 Summary data

Supplementary Figure 1 / Supplementary Table 5 Summary data

Supplementary Table 6 Summary data

Supplementary Figure 2 Summary data

Supplementary Figure 3 Summary data

## Acknowledgements

We thank Marius Moisa, Karl Treiber, Jennifer Hueber, Rosa Bohlender, Athina Tzovara and Filip Melinščak for assistance. This work was funded by Swiss National Foundation grant 320030_149586/1 to DRB and Wellcome Centre for Human Neuroimaging (University College London, UK) core funding 203147/Z/16/Z. DRB and CCR are supported by funding from the European Research Council (ERC) under the European Union’s Horizon 2020 research and innovation programme (Grant agreements No. ERC-2018 CoG-816564 ActionContraThreat to DRB; ERC-2016 CoG-725355 BRAINCODES to CCR). DRB receives support from the National Institute for Health Research (NIHR) UCLH Biomedical Research Centre. CCR receives support from the Swiss National Science Foundation (100019L_173248).

## Author contributions

DRB, MS and CCR designed the experiment; CCR provided methodological expertise, training, and access to equipment; MS piloted the experiment; KEO and MS acquired the data; KEO analyzed the data; KEO wrote the original draft of the article; MS, CCR and DRB contributed to the editing and revising of the paper; DRB supervised work at all stages.

## Code and data availability statement

The code for the psychophysiological analyses presented in the manuscript is available publicly on GitLab: https://gitlab.com/kojala/tms_somatosensory_fc. The full anonymized data set (except for individual MRI/fMRI data) and fMRI analysis code will be available at publication of the article or at request from KEO. Summary data to reproduce the figures and tables in the manuscript and supplementary information are published together with this manuscript.

## Materials & correspondence

Correspondence should be addressed to DRB and KEO.

## References

1. Duits, P. et al. Updated meta-analysis of classical fear conditioning in the anxiety disorders. Depress. Anxiety 32, 239–253 (2015).

2. Dunsmoor, J. E. & Paz, R. Fear Generalization and Anxiety: Behavioral and Neural Mechanisms. Biol. Psychiatry 78, 336–343 (2015).

3. Pearce, J. M. & Bouton, M. E. Theories of Associative Learning in Animals. Annu. Rev. Psychol. 52, 111–139 (2001).

4. Yau, J. O. Y. & McNally, G. P. Brain Mechanisms Controlling Pavlovian Fear Conditioning. J. Exp. Psychol. Anim. Learn. Cogn. 44, 341–357 (2018).

5. Flores, Á., Fullana, M., Soriano-Mas, C. & Andero, R. Lost in translation: how to upgrade fear memory research. Mol. Psychiatry 1–11 (2018) doi:10.1038/s41380-017-0006-0.

6. Phelps, E. A. & Hofmann, S. G. Memory editing from science fiction to clinical practice. Nature 572, 43–50 (2019).

7. Maren, S. & Quirk, G. J. Neuronal signalling of fear memory. Nat. Neurosci. Rev. 5, 844–852 (2004).

8. Tovote, P., Fadok, J. P. & Lüthi, A. Neuronal circuits for fear and anxiety. Nat. Rev. Neurosci. 16, 317–331 (2015).

9. McDonald, A. J. Cortical pathways to the mammalian amygdala. Prog. Neurobiol. 55, 257–332 (1998).

10. Amaral, D. G., Behniea, H. & Kelly, J. L. Topographic organization of projections from the amygdala to the visual cortex in the macaque monkey. Neuroscience 118, 1099–1120 (2003).

11. Abivardi, A. & Bach, D. R. Deconstructing white matter connectivity of human amygdala nuclei with thalamus and cortex subdivisions in vivo. Hum. Brain Mapp. 38, 3927–3940 (2017).

12. Sacco, T. & Sacchetti, B. Role of Secondary Sensory Cortices. Science (80-.). 329, 649–657 (2010).

13. Galván, V. V. & Weinberger, N. M. Long-term consolidation and retention of learning-induced tuning plasticity in the auditory cortex of the guinea pig. Neurobiol. Learn. Mem. 77, 78–108 (2002).

14. Campeau, S. & Davis, M. Involvement of subcortical and cortical afferents to the lateral nucleus of the amygdala in fear conditioning measured with fear-potentiated startle in rats trained concurrently with auditory and visual conditioned stimuli. J. Neurosci. 15, 2312–2327 (1995).

15. Romanski, L. M. & LeDoux, J. E. Bilateral destruction of neocortical and perirhinal projection targets of the acoustic thalamus does not disrupt auditory fear conditioning. Neurosci. Lett. 142, 228–232 (1992).

16. Romanski, L. M. & LeDoux, J. E. Equipotentiality of thalamo-amygdala and thalamo-cortico-amygdala circuits in auditory fear conditioning. J. Neurosci. 12, 4501–4509 (1992).

17. Peter, M. et al. Induction of immediate early genes in the mouse auditory cortex after auditory cued fear conditioning to complex sounds. Genes, Brain Behav. 11, 314–324 (2012).

18. Moczulska, K. E. et al. Dynamics of dendritic spines in the mouse auditory cortex during memory formation and memory recall. Proc. Natl. Acad. Sci. U. S. A. 110, 18315–18320 (2013).

19. Banerjee, S. B. et al. Perineuronal Nets in the Adult Sensory Cortex Are Necessary for Fear Learning. Neuron 95, 169–179.e3 (2017).

20. Gillet, S. N., Kato, H. K., Justen, M. A., Lai, M. & Isaacson, J. S. Fear learning regulates cortical sensory representations by suppressing habituation. Front. Neural Circuits 11, 1–9 (2018).

21. Yang, Y. et al. Selective synaptic remodeling of amygdalocortical connections associated with fear memory. Nat. Neurosci. 19, 1348–1355 (2016).

22. Dalmay, T. et al. A critical role for neocortical processing of threat memory. Neuron 104, 1180–1194.e7 (2019).

23. Wigestrand, M. B., Schiff, H. C., Fyhn, M., LeDoux, J. E. & Sears, R. M. Primary auditory cortex regulates threat memory specificity. Learn. Mem. 24, 55–58 (2017).

24. Antunes, R. & Moita, M. A. Discriminative auditory fear learning requires both tuned and nontuned auditory pathways to the amygdala. J. Neurosci. 30, 9782–9787 (2010).

25. Aizenberg, M., Mwilambwe-Tshilobo, L., Briguglio, J. J., Natan, R. G. & Geffen, M. N. Bidirectional Regulation of Innate and Learned Behaviors That Rely on Frequency Discrimination by Cortical Inhibitory Neurons. PLoS Biol. 13, 1–32 (2015).

26. Letzkus, J. J. et al. A disinhibitory microcircuit for associative fear learning in the auditory cortex. Nature 480, 331–335 (2011).

27. Kraus, M. et al. Memory consolidation for the discrimination of frequency-modulated tones in Mongolian gerbils is sensitive to protein-synthesis inhibitors applied to the auditory cortex. Learn. Mem. 9, 293–303 (2002).

28. Zhang, G.-W. et al. A Non-canonical Reticular-Limbic Central Auditory Pathway via Medial Septum Contributes to Fear Conditioning. Neuron 97, 406–417.e4 (2018).

29. Keil, A., Stolarova, M., Moratti, S. & Ray, W. J. Adaptation in human visual cortex as a mechanism for rapid discrimination of aversive stimuli. Neuroimage 36, 472–479 (2007).

30. Miskovic, V. & Keil, A. Perceiving Threat In the Face of Safety: Excitation and Inhibition of Conditioned Fear in Human Visual Cortex. J. Neurosci. 33, 72–78 (2013).

31. Petro, N. M. et al. Multimodal Imaging Evidence for a Frontocortical Modulation of Visual Cortex during the Selective Processing of Conditioned Threat. J. Cogn. Neurosci. 29, 953/967 (2017).

32. Li, W., Howard, J. D., Parrish, T. B. & Gottfried, J. A. Aversive learning enhances perceptual and cortical discrimination of indiscriminable odor cues. Science (80-.). 319, 1842–1845 (2008).

33. Apergis-Schoute, A. M., Schiller, D., LeDoux, J. E. & Phelps, E. A. Extinction resistant changes in the human auditory association cortex following threat learning. Neurobiol. Learn. Mem. 113, 109–114 (2014).

34. Staib, M. & Bach, D. R. Stimulus-invariant auditory cortex threat encoding during fear conditioning with simple and complex sounds. Neuroimage 166, 276–284 (2018).

35. Staib, M., Abivardi, A. & Bach, D. R. Primary auditory cortex representation of fear-conditioned musical sounds. Hum. Brain Mapp. 1–10 (2019) doi:10.1002/hbm.24846.

36. You, Y., Brown, J. & Li, W. Human Sensory Cortex Contributes to the Long-Term Storage of Aversive Conditioning. J. Neurosci. 41, 3222–3233 (2021).

37. Huang, Y. Z., Edwards, M. J., Rounis, E., Bhatia, K. P. & Rothwell, J. C. Theta burst stimulation of the human motor cortex. Neuron 45, 201–206 (2005).

38. Gazzaniga, M. S. Cerebral specialization and interhemispheric communication. Does the corpus callosum enable the human condition? Brain 123, 1293–1326 (2000).

39. Harvie, D. S. et al. When touch predicts pain: Predictive tactile cues modulate perceived intensity of painful stimulation independent of expectancy. Scand. J. Pain 11, 11–18 (2016).

40. Harvie, D. S. et al. Selectivity of conditioned fear of touch is modulated by somatosensory precision. Psychophysiology 53, 921–929 (2016).

41. Korn, C. W., Staib, M., Tzovara, A., Castegnetti, G. & Bach, D. R. A pupil size response model to assess fear learning. Psychophysiology 54, 330–343 (2017).

42. Polanía, R., Nitsche, M. A. & Ruff, C. C. Studying and modifying brain function with non-invasive brain stimulation. Nat. Neurosci. 21, (2018).

43. Siebner, H. R. & Rothwell, J. Transcranial magnetic stimulation: New insights into representational cortical plasticity. Exp. Brain Res. 148, 1–16 (2003).

44. Huang, Y. Z., Chen, R. S., Rothwell, J. C. & Wen, H. Y. The after-effect of human theta burst stimulation is NMDA receptor dependent. Clin. Neurophysiol. 118, 1028–1032 (2007).

45. Li, C. T. et al. Critical role of glutamatergic and GABAergic neurotransmission in the central mechanisms of theta-burst stimulation. Hum. Brain Mapp. 40, 2001–2009 (2019).

46. Stagg, C. J. et al. Neurochemical effects of theta burst stimulation as assessed by magnetic resonance spectroscopy. J. Neurophysiol. 101, 2872–2877 (2009).

47. Morey, R. D. Confidence Intervals from Normalized Data: A correction to Cousineau (2005). Tutor. Quant. Methods Psychol. 4, 61–64 (2008).

48. Baltaci, S. B., Mogulkoc, R. & Baltaci, A. K. Molecular Mechanisms of Early and Late LTP. Neurochem. Res. 44, 281–296 (2019).

49. Mulckhuyse, M., Engelmann, J. B., Schutter, D. J. L. G. & Roelofs, K. Right posterior parietal cortex is involved in disengaging from threat: A 1-Hz rTMS study. Soc. Cogn. Affect. Neurosci. 12, 1814–1822 (2017).

50. Borgomaneri, S. et al. State-Dependent TMS over Prefrontal Cortex Disrupts Fear-Memory Reconsolidation and Prevents the Return of Fear. Curr. Biol. 1–8 (2020) doi:10.1016/j.cub.2020.06.091.

51. Herrmann, M. J., Mühlberger, A., Ehlis, A. C., Deckert, J. & Polak, T. A systematic review of non-invasive brain stimulation and fear extinction. PsyArXiv May 24 (2019) doi:https://doi.org/10.31234/osf.io/u65va.

52. Baek, K., Chae, J. H. & Jeong, J. The effect of repetitive transcranial magnetic stimulation on fear extinction in rats. Neuroscience 200, 159–165 (2012).

53. Guhn, A. et al. Medial prefrontal cortex stimulation modulates the processing of conditioned fear. Front. Behav. Neurosci. 8, 1–11 (2014).

54. Raij, T. et al. Prefrontal Cortex Stimulation Enhances Fear Extinction Memory in Humans. Biol. Psychiatry 84, 129–137 (2018).

55. Hamm, A. O. et al. Affective blindsight: Intact fear conditioning to a visual cue in a cortically blind patient. Brain 126, 267–275 (2003).

56. Badura-Brack, A. S. et al. Decreased somatosensory activity to non-threatening touch in combat veterans with posttraumatic stress disorder. Psychiatry Res. - Neuroimaging 233, 194–200 (2015).

57. Gdalyahu, A. et al. Associative fear learning enhances sparse network coding in primary sensory cortex. Neuron 75, 121–132 (2012).

58. Joachimsthaler, B., Brugger, D., Skodras, A. & Schwarz, C. Spine loss in primary somatosensory cortex during trace eyeblink conditioning. J. Neurosci. 35, 3772–3781 (2015).

59. Tokarski, K., Urban-Ciecko, J., Kossut, M. & Hess, G. Sensory learning-induced enhancement of inhibitory synaptic transmission in the barrel cortex of the mouse. Eur. J. Neurosci. 26, 134–141 (2007).

60. Weinberger, N. M. Associative representational plasticity in the auditory cortex: Resolving conceptual and empirical problems. Learn. Mem. 14, 1–16 (2007).

61. Kholodar-Smith, D. B., Allen, T. A. & Brown, T. H. Fear Conditioning to Discontinuous Auditory Cues Requires Perirhinal Cortical Function. Behav. Neurosci. 122, 1178–1185 (2008).

62. Bach, D. R., Tzovara, A. & Vunder, J. Blocking human fear memory with the matrix metalloproteinase inhibitor doxycycline. Mol. Psychiatry 23, 1584–1589 (2018).

63. Bach, D. R., Melinščak, F., Fleming, S. M. & Voelkle, M. C. Calibrating the experimental measurement of psychological attributes. Nat. Hum. Behav. 4, 1229–1235 (2020).

64. Bestmann, S. & Feredoes, E. Combined neurostimulation and neuroimaging in cognitive neuroscience: Past, present, and future. Ann. N. Y. Acad. Sci. 1296, 11–30 (2013).

65. Valchev, N. et al. cTBS delivered to the left somatosensory cortex changes its functional connectivity during rest. Neuroimage 114, 386–397 (2015).

66. Vlachos, A. et al. Repetitive magnetic stimulation induces functional and structural plasticity of excitatory postsynapses in mouse organotypic hippocampal slice cultures. J. Neurosci. 32, 17514–17523 (2012).

67. Nitsche, M. A., Müller-Dahlhaus, F., Paulus, W. & Ziemann, U. The pharmacology of neuroplasticity induced by non-invasive brain stimulation: Building models for the clinical use of CNS active drugs. J. Physiol. 590, 4641–4662 (2012).

68. Fritsch, B. et al. Direct current stimulation promotes BDNF-dependent synaptic plasticity: Potential implications for motor learning. Neuron 66, 198–204 (2010).

69. Cheeran, B. et al. A common polymorphism in the brain-derived neurotrophic factor gene (BDNF) modulates human cortical plasticity and the response to rTMS. J. Physiol. 586, 5717–5725 (2008).

70. Ueyama, E. et al. Chronic repetitive transcranial magnetic stimulation increases hippocampal neurogenesis in rats. Psychiatry Clin. Neurosci. 65, 77–81 (2011).

71. Suppa, A. et al. Ten Years of Theta Burst Stimulation in Humans: Established Knowledge, Unknowns and Prospects. Brain Stimul. 9, 323–335 (2016).

72. Abraham, W. C. & Huggett, A. Induction and reversal of long-term potentiation by repeated high-frequency stimulation in rat hippocampal slices. Hippocampus 7, 137–145 (1997).

73. Shouval, H. Z., Bear, M. F. & Cooper, L. N. A unified model of NMDA receptor-dependent bidirectional synaptic plasticity. Proc. Natl. Acad. Sci. U. S. A. 99, 10831–10836 (2002).

74. Silvanto, J., Bona, S., Marelli, M. & Cattaneo, Z. On the mechanisms of Transcranial Magnetic Stimulation (TMS): How brain state and baseline performance level determine behavioral effects of TMS. Front. Psychol. 9, 1–8 (2018).

75. Kroes, M. C. W., Schiller, D., LeDoux, J. E. & Phelps, E. A. Translational Approaches Targeting Reconsolidation. Curr. Top. Behav. Neurosci. 28, 197–230 (2016).

76. Brainard, D. H. The Psychophysics Toolbox. Spat. Vis. 10, 433–436 (1997).

77. Sladky, R. et al. Slice-timing effects and their correction in functional MRI. Neuroimage 58, 588–594 (2011).

78. Ashburner, J. & Friston, K. J. Unified segmentation. Neuroimage 26, 839–851 (2005).

79. Martuzzi, R., van der Zwaag, W., Farthouat, J., Gruetter, R. & Blanke, O. Human finger somatotopy in areas 3b, 1, and 2: A 7T fMRI study using a natural stimulus. Hum. Brain Mapp. 35, 213–226 (2014).

80. Stokes, M. G. et al. Distance-adjusted motor threshold for transcranial magnetic stimulation. Clin. Neurophysiol. 118, 1617–1625 (2007).

81. Bach, D. R. et al. Psychophysiological modeling: Current state and future directions. Psychophysiology 55, e13209 (2018).

82. Bach, D. R., Daunizeau, J., Friston, K. J. & Dolan, R. J. Dynamic causal modelling of anticipatory skin conductance responses. Biol. Psychol. 85, 163–170 (2010).

83. Staib, M., Castegnetti, G. & Bach, D. R. Optimising a model-based approach to inferring fear learning from skin conductance responses. J. Neurosci. Methods 255, 131–138 (2015).

84. Bach, D. R., Flandin, G., Friston, K. J. & Dolan, R. J. Time-series analysis for rapid event-related skin conductance responses. J. Neurosci. Methods 184, 224–234 (2009).

85. Bach, D. R., Flandin, G., Friston, K. J. & Dolan, R. J. Modelling event-related skin conductance responses. Int. J. Psychophysiol. 75, 349–356 (2010).

86. Gerster, S., Namer, B., Elam, M. & Bach, D. R. Testing a linear time invariant model for skin conductance responses by intraneural recording and stimulation. Psychophysiology 55, (2018).

87. Korn, C. W. & Bach, D. R. A solid frame for the window on cognition: Modeling event-related pupil responses. J. Vis. 16, 28 (2016).

88. Sjouwerman, R., Niehaus, J., Kuhn, M. & Lonsdorf, T. B. Don’t startle me—Interference of startle probe presentations and intermittent ratings with fear acquisition. Psychophysiology 53, 1889–1899 (2016).

89. Blumenthal, T. D. et al. Committee report: Guidelines for human startle eyeblink electromyographic studies. Psychophysiology 42, 1–15 (2005).

90. Khemka, S., Tzovara, A., Gerster, S., Quednow, B. & Bach, D. R. Modelling startle eye-blink electromyogram to assess fear learning. Psychophysiology 54, 204–214 (2017).

91. RCoreTeam. R: A Language and Environment for Statistical Computing. (2020).

92. RStudioTeam. RStudio: Integrated Development for R. (2020).

93. Bach, D. R., Näf, M., Deutschmann, M., Tyagarajan, S. K. & Quednow, B. B. Threat memory reminder under matrix metalloproteinase 9 inhibitor doxycycline globally reduces subsequent memory plasticity. J. Neurosci. 39, 1285–19 (2019).

94. Lenth, R. V et al. emmeans: Estimated marginal means. (2021).

95. Ben-Shachar, M., Lüdecke, D. & Makowski, D. effectsize: Estimation of Effect Size Indices and Standardized Parameters. J. Open Source Softw. 5, 2815 (2020).

96. Pinheiro, J., Bates, D., DebRoy, S., Sarkar, D. & RCoreTeam. nlme: Linear and Nonlinear Mixed Effects Models. (2020).

97. Makowski, D., Ben-Shachar, M. & Lüdecke, D. bayestestR: Describing Effects and their Uncertainty, Existence and Significance within the Bayesian Framework. J. Open Source Softw. 4, 1541 (2019).

98. Wagenmakers, E. J. A practical solution to the pervasive problems of p values. Psychon. Bull. Rev. 14, 779–804 (2007).

