## Supplementary material for "Transcranial theta-burst stimulation of primary sensory cortex attenuates somatosensory threat memory in humans": Ojala et al. 2021 Supplementary Information

**Supplementary Table 1. Statistical test results for pairwise comparisons of condition-wise startle eye-blink responses (SEBR) during threat memory retention test on test day 2.**

| Paired t-tests | CS | df | *t* | *p* | *D* |
|---|---|---|---|---|---|
| Control group CS+ > CS− | Simple | 26 | 3.99 | < 0.001 | 0.767 |
| | Complex | 26 | 1.52 | 0.07 | 0.292 |
| Experimental group CS+ > CS− | Simple | 24 | −0.78 | 0.78 | −0.157 |
| | Complex | 24 | 0.36 | 0.36 | 0.072 |

*df* = Degrees of Freedom, *d* = Cohen's *d*. P-values are uncorrected for multiple comparisons.

**Supplementary Table 2. Statistical test results for trial-wise startle eye-blink responses (SEBR) during threat memory retention test on test day 2.**

| Linear mixed effects model | df1 | df2 | *F* | *p* |
|---|---|---|---|---|
| Group | 1 | 50 | 5.71 | 0.021 |
| Trial | 1 | 1182 | 180.6 | < .0001 |
| CS type | 1 | 1182 | 3.83 | 0.051 |
| CS complexity | 1 | 1182 | 0.72 | 0.40 |
| Group x Trial | 1 | 1182 | 0.04 | 0.84 |
| Group x CS type | 1 | 1182 | 5.49 | 0.019 |
| Trial x CS type | 1 | 1182 | 1.70 | 0.19 |
| Group x CS complexity | 1 | 1182 | 0.11 | 0.74 |
| Trial x CS complexity | 1 | 1182 | 12.6 | < 0.001 |
| CS type x CS complexity | 1 | 1182 | 0.01 | 0.93 |
| Group x Trial x CS type | 1 | 1182 | 0.97 | 0.32 |
| Group x Trial x CS complexity | 1 | 1182 | 0.27 | 0.60 |
| Group x CS type x CS complexity | 1 | 1182 | 2.57 | 0.11 |
| Trial x CS type x CS complexity | 1 | 1182 | 2.55 | 0.11 |
| Group x Trial x CS type x CS complexity | 1 | 1182 | 0.63 | 0.43 |

df = Degrees of Freedom, *d* = Cohen's *d*. Model structure with nlme in R: SEBR ~ Group * Trial * CS type * CS complexity + (CS complexity * Trial)|Subject.

**Supplementary Table 3. Statistical test results for pairwise comparisons of skin conductance responses (SCR) and pupil size responses (PSR) during threat conditioning on test day 1 directly after theta-burst TMS.**

| Paired t-tests for SCR | CS | df | *t* | *p* | *D* |
|---|---|---|---|---|---|
| Control group CS+ > CS− | Simple | 33 | 2.10 | 0.022 | 0.36 |
| | Complex | 33 | 0.30 | 0.38 | 0.05 |
| Experimental group CS+ > CS− | Simple | 27 | 2.97 | 0.003 | 0.56 |
| | Complex | 27 | 1.49 | 0.074 | 0.28 |
| Paired t-tests for PSR | CS | df | *t* | *p* | *d* |
| Control group CS+/CS− | Simple | 16 | 1.72 | 0.053 | 0.42 |
| | Complex | 16 | 2.31 | 0.017 | 0.56 |
| Experimental group CS+/CS− | Simple | 19 | 0.37 | 0.36 | 0.08 |
| | Complex | 19 | 0.95 | 0.18 | 0.21 |

*df* = Degrees of Freedom, *d* = Cohen's *d*. P-values are uncorrected for multiple comparisons.

**Supplementary Table 4. Statistical test results for trial-wise skin conductance responses (SCR) during threat conditioning on test day 1 directly after theta-burst TMS.**

| Linear mixed effects model | df1 | df2 | *F* | *p* |
|---|---|---|---|---|
| Group | 1 | 60 | 0.32 | 0.57 |
| Trial | 1 | 4310 | 232.2 | < 0.0001 |
| CS type | 1 | 4310 | 16.6 | < 0.0001 |
| CS complexity | 1 | 4310 | 7.04 | 0.008 |
| Group x Trial | 1 | 4310 | 0.76 | 0.38 |
| Group x CS type | 1 | 4310 | 1.63 | 0.20 |
| Trial x CS type | 1 | 4310 | 18.8 | < 0.0001 |
| Group x CS complexity | 1 | 4310 | 0.27 | 0.60 |
| Trial x CS complexity | 1 | 4310 | 26.9 | < 0.0001 |
| CS type x CS complexity | 1 | 4310 | 14.5 | 0.0001 |
| Group x Trial x CS type | 1 | 4310 | 0.80 | 0.37 |
| Group x Trial x CS complexity | 1 | 4310 | 5.67 | 0.017 |
| Group x CS type x CS complexity | 1 | 4310 | 2.86 | 0.091 |
| Trial x CS type x CS complexity | 1 | 4310 | 3.91 | 0.048 |
| Group x Trial x CS type x CS complexity | 1 | 4310 | 0.11 | 0.74 |

df = Degrees of Freedom, *d* = Cohen's *d*. Model structure with nlme in R: SCR ~ Group * Trial * CS type * CS complexity + 1|Subject.

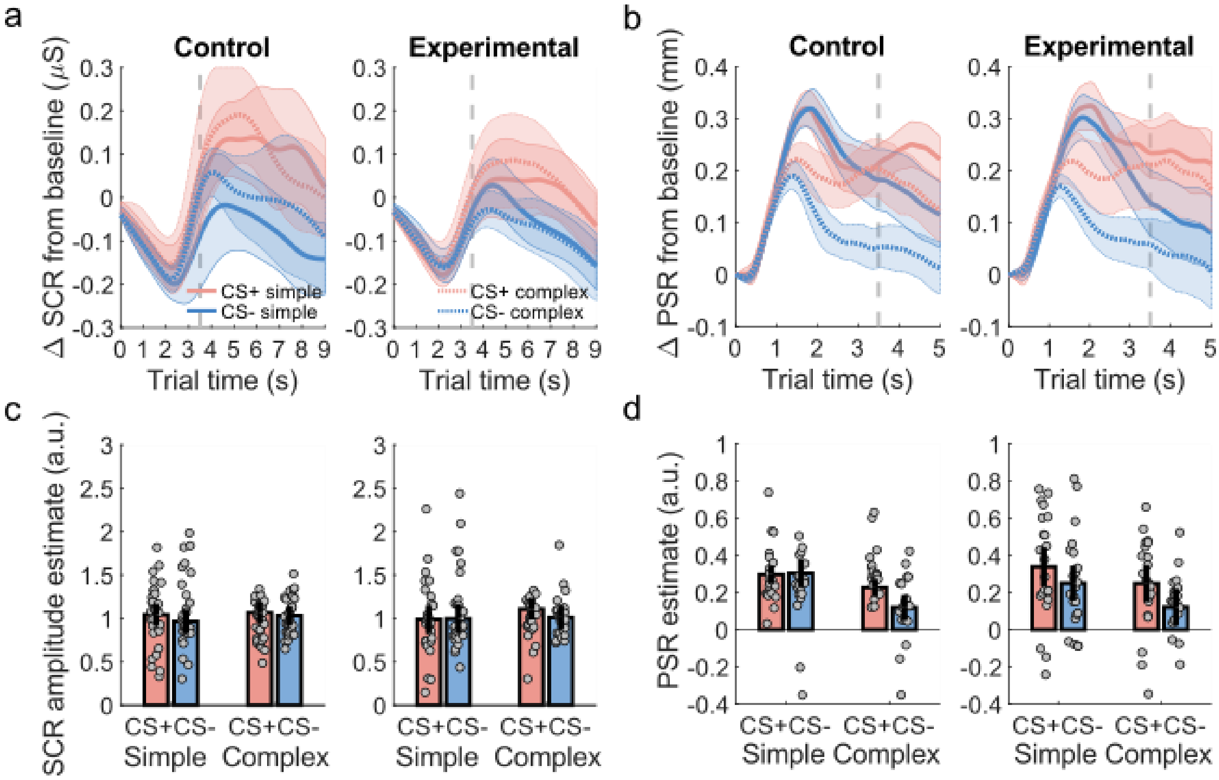


**Supplementary Figure 1.** Skin conductance responses (SCR) and pupil size responses (PSR) during threat relearning on test day 2. **a-b**. Averaged trial time courses of skin conductance and pupil size changes from baseline for the different stimuli, separately for control and experimental groups. Shaded areas represent within-subject standard errors of the mean; dashed grey line marks US onset (only no-US trials were included); $\mu$S are micro-Siemens. **b-c**. PSR estimates showed that the participants learned the CS+/CS− difference overall (main effect of CS type, Supplementary Table 3), evidencing threat conditioning. There was no evidence for a difference in relearning between the control and experimental groups. PSR for simple stimuli were on average larger than for complex stimuli, but this was not different for the two groups or between CS+ and CS−. Error bars show 95% within-subject standard errors of the mean reflecting paired, one-tailed CS+ > CS− comparison; a. u. are arbitrary units.

**Supplementary Table 5. Statistical test results for condition-wise skin conductance and pupil size responses during threat re-learning on test day 2 after threat memory retention test.**

| Repeated-measures ANOVA for SCR | | df1 | df2 | *F* | *p* | $\eta^2$ |
|---|---|---|---|---|---|---|
| Group | | 1 | 54 | 0 | 0.98 | < 0.001 |
| CS type | | 1 | 54 | 1.44 | 0.24 | 0.006 |
| CS complexity | | 1 | 54 | 0.79 | 0.38 | 0.007 |
| Group x CS type | | 1 | 54 | 0.02 | 0.88 | < 0.001 |
| Group x CS complexity | | 1 | 54 | 0.03 | 0.86 | < 0.001 |
| CS type x CS complexity | | 1 | 54 | 0.26 | 0.61 | < 0.001 |
| Group x CS type x CS complexity | | 1 | 54 | 1.44 | 0.24 | 0.003 |
| Paired t-tests for SCR | CS | df | | *t* | *p* | *d* |
| Control group CS+ > CS− | Simple | 28 | | 1.22 | 0.12 | 0.23 |
| | Complex | 28 | | 0.56 | 0.29 | 0.10 |
| Experimental group CS+ > CS− | Simple | 26 | | −0.11 | 0.54 | 0.02 |
| | Complex | 26 | | 1.07 | 0.15 | 0.21 |

| Repeated-measures ANOVA for PSR | | df1 | df2 | *F* | *p* | $\eta^2$ |
|---|---|---|---|---|---|---|
| Group | | 1 | 41 | 0.091 | 0.77 | < 0.001 |
| CS type | | 1 | 41 | 7.04 | 0.011 | 0.036 |
| CS complexity | | 1 | 41 | 24.10 | < 0.001 | 0.075 |
| Group x CS type | | 1 | 41 | 0.99 | 0.33 | 0.006 |
| Group x CS complexity | | 1 | 41 | 0.012 | 0.92 | < 0.001 |
| CS type x CS complexity | | 1 | 41 | 1.66 | 0.20 | 0.006 |
| Group x CS type x CS complexity | | 1 | 41 | 0.69 | 0.41 | 0.002 |
| Paired t-tests for PSR | CS | df | | *t* | *p* | *d* |
| Control group CS+ > CS− | Simple | 20 | | −0.18 | 0.57 | −0.040 |
| | Complex | 20 | | 2.16 | 0.022 | 0.471 |
| Experimental group CS+ > CS− | Simple | 21 | | 1.42 | 0.086 | 0.302 |
| | Complex | 21 | | 1.82 | 0.041 | 0.388 |

df = Degrees of Freedom, $\eta^2$ = Eta squared (explained variance), *d* = Cohen's *d*. P-values are uncorrected for multiple comparisons.

**Supplementary Table 6. Statistical test results for trial-wise skin conductance responses (SCR) during threat re-learning on test day 2 after threat memory retention test.**

| Linear mixed effects model | df1 | df2 | *F* | *p* |
|---|---|---|---|---|
| Group | 1 | 54 | 0.001 | 0.97 |
| Trial | 1 | 3886 | 67.8 | < 0.0001 |
| CS type | 1 | 3886 | 5.83 | 0.016 |
| CS complexity | 1 | 3886 | 2.10 | 0.15 |
| Group x Trial | 1 | 3886 | 8.91 | 0.003 |
| Group x CS type | 1 | 3886 | 2.13 | 0.15 |
| Trial x CS type | 1 | 3886 | 6.55 | 0.011 |
| Group x CS complexity | 1 | 3886 | < 0.001 | 0.99 |
| Trial x CS complexity | 1 | 3886 | 22.8 | < 0.0001 |
| CS type x CS complexity | 1 | 3886 | 3.35 | 0.067 |
| Group x Trial x CS type | 1 | 3886 | 3.42 | 0.065 |
| Group x Trial x CS complexity | 1 | 3886 | 0.009 | 0.93 |
| Group x CS type x CS complexity | 1 | 3886 | 1.31 | 0.25 |
| Trial x CS type x CS complexity | 1 | 3886 | 2.97 | 0.085 |
| Group x Trial x CS type x CS complexity | 1 | 3886 | 1.33 | 0.25 |

df = Degrees of Freedom, *d* = Cohen's *d*. Model structure with nlme in R: SCR ~ Group * Trial * CS type * CS complexity + 1|Subject.

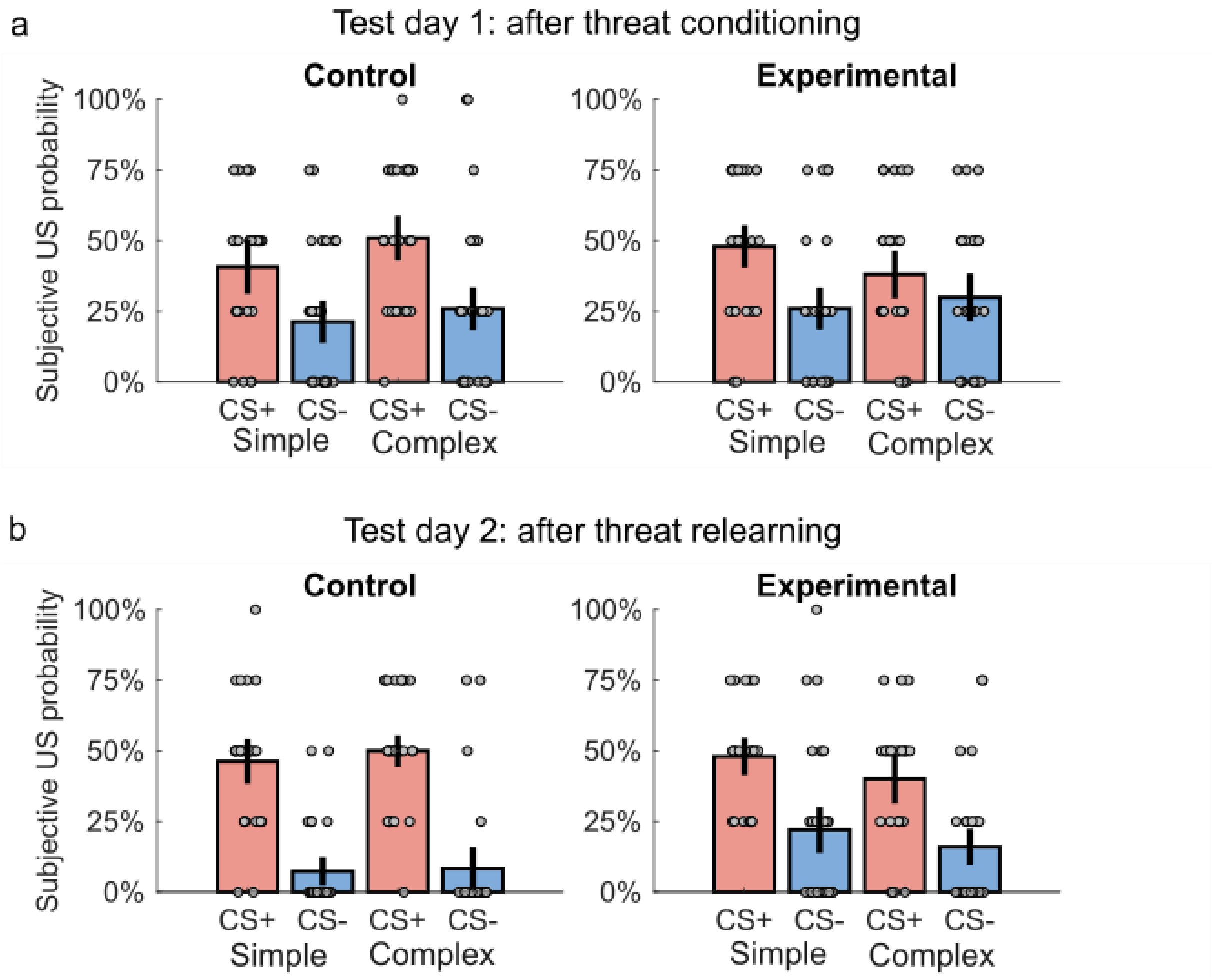


**Supplementary Figure 2.** CS-US contingency awareness as measured by subjective US probability rating with possible levels 0%, 25%, 50%, 75% and 100%. **a**. Overall, participants explicitly learned to differentiate CS+ and CS− on test day 1. Experimental group explicitly learned the CS+/CS− difference for simple but not for complex stimuli. **b**. Participants learned to differentiate the CSs also on test day 2. Qualitatively, it seems that the participants' explicit learning improved on test day 2 from test day 1. However, experimental group learned to differentiate CS+ and CS− less well than the control group. Error bars show 95% within-subject standard errors of the mean reflecting the paired, one-tailed CS+ > CS− comparison.

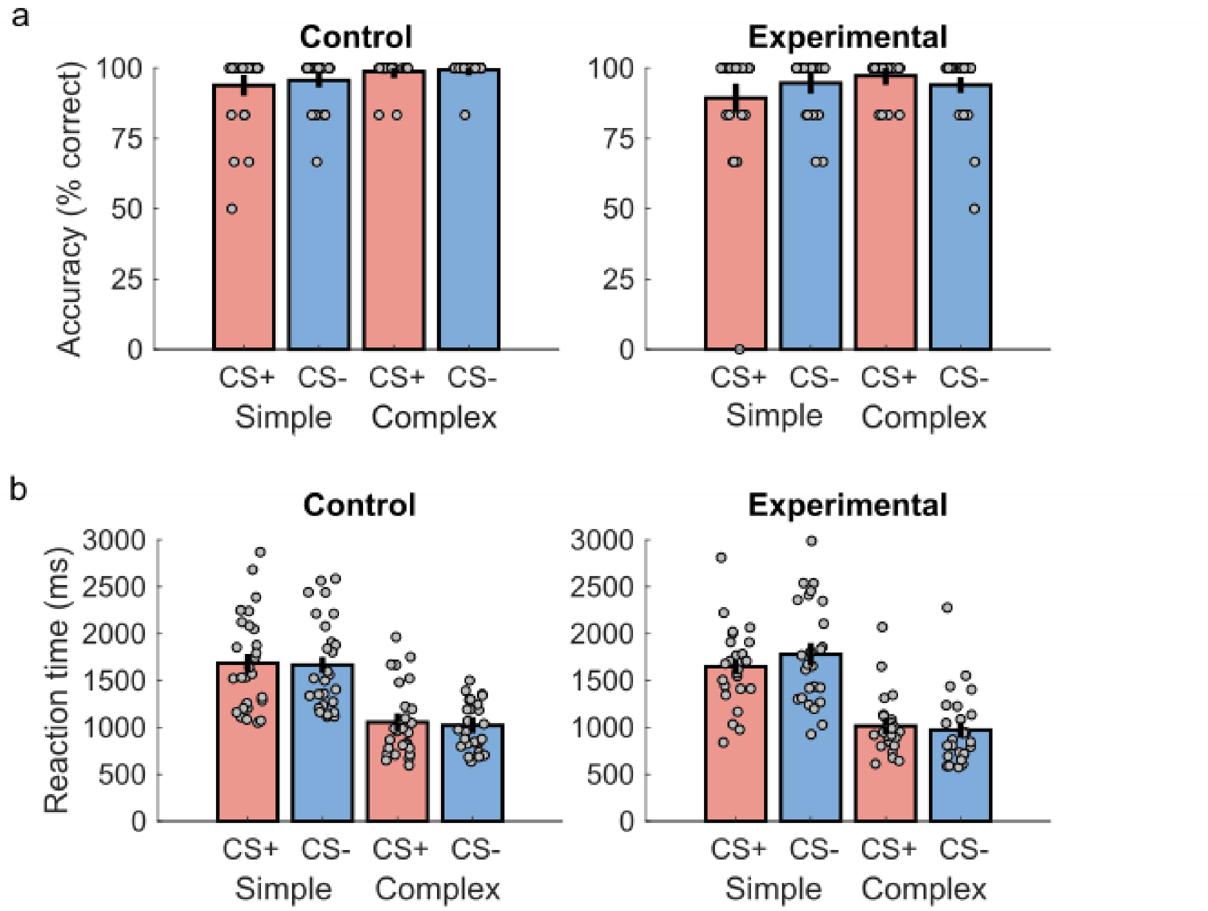


**Supplementary Figure 3.** Accuracy (% correct responses) and reaction times during threat memory retention test on test day 2. **a**. Participants were on average more accurate on complex than on simple CS trials. **b**. Participants were on average faster to respond to complex than simple CS. There were no significant differences between groups nor CS+/CS− in either accuracy or reaction times. Error bars show 95% within-subject standard errors of the mean reflecting the paired, one-tailed CS+ > CS− comparison.
